# *Drosophila melanogaster* establishes a species-specific mutualistic interaction with stable gut-colonizing bacteria

**DOI:** 10.1101/265991

**Authors:** Inês S. Pais, Rita S. Valente, Marta Sporniak, Luis Teixeira

## Abstract

Animals live together with diverse bacteria that can impact their biology. In *Drosophila melanogaster*, gut-associated bacterial communities are relatively simple in composition but also have a strong impact on host development and physiology. However, it is still unknown if bacteria can proliferate and stably associate with the gut of *D. melanogaster*. In fact, it is generally assumed that bacteria are transient and their constant ingestion with food is required to maintain their presence in the gut. Here, we identify bacterial species from wild-caught *D. melanogaster* that stably associate with the host independently of continuous inoculation. Moreover, we show that specific *Acetobacter* wild isolates can proliferate in the gut. We further demonstrate that the interaction between *D. melanogaster* and the wild isolated *Acetobacter thailandicus* is mutually beneficial and that the stability of the gut association is key to this mutualism. The stable population in the gut of *D. melanogaster* allows continuous bacterial spreading into the environment, which is advantageous to the bacterium itself. The bacterial dissemination is in turn advantageous to the host since the next generation of flies develops in the presence of this particularly beneficial bacterium. *Ac. thailandicus* leads to a faster host development and higher fertility of emerging adults, when compared to other bacteria isolated from wild-caught flies. Furthermore, *Ac. thailandicus* is sufficient and advantageous when *D. melanogaster* develops in axenic or freshly collected figs, respectively. This isolate of *Ac. thailandicus* colonizes several genotypes of *D. melanogaster* but not of the closely related *Drosophila simulans*, indicating that the stable association is host specific. This work establishes a new conceptual model to understand *D. melanogaster*- gut microbiota interactions in an ecological context; stable interactions can be mutualistic through microbial farming, a common strategy in insects. Moreover, these results develop the use of *D. melanogaster* as a model to study gut microbiota proliferation and colonization.

**Author summary:** Animals, including humans, live together with complex bacterial communities in their gut that influence their physiology and health. The fruit fly *Drosophila melanogaster* has been an excellent model organism to study host-microbe interactions and harbours a relative simple gut bacterial community. However, it is not known which of these bacteria can proliferate and form stable communities in the gut, and the current hypothesis is that these bacteria are only transiently associated with the gut. Here, we show that in *D. melanogaster* collected from a natural population stable gut bacteria do exist. We isolated specific species that can proliferate in the gut and form a stable association. This is beneficial to the bacteria since they can be constantly spread by the flies as they move around. On the other hand, this is a form of farming as the next generation of flies benefit from the association with these particular bacteria during development. They become adults faster and are more fertile than if they develop with other bacteria encountered in nature. These advantages are also observed when flies develop in figs, a natural food source. Our findings show that *D. melanogaster* has stable colonizing bacteria in the gut and establish a new framework to study host-gut bacteria interactions.

## Introduction

Animals live with microbial communities that have a strong impact on their physiology, including their development, nutrition, immunity and behavior [1]. These effects may be partially explained by adaptation of animals to the ubiquitous presence of microbes and integration of this cue in their developmental and physiological programs. However, association with specific microbes may increase their fitness in the environment they live or provide the capacity to explore new niches. For instance, many endosymbionts in insects provide essential metabolites, allowing hosts to explore food sources deficient in some nutrients, as plant sap and blood [2-6].

A primary organ for animal-microbe interactions is the gut, which is an interface between the external environment and the animal body. The gut microbiota can be very complex and comprised of up to one thousand different bacterial species, as in humans [7]. Its composition varies to different degrees between and within host species. Moreover, even within the same host it can be very dynamic and fluctuate with host age and health, diet, and other environmental conditions [8-11]. Understanding the composition of the gut microbiota, which factors regulate it, and how these interactions impact both the host and the microbes are, therefore, major research questions.

*Drosophila melanogaster* has been used as model system to study host interaction with gut bacteria [12,13]. Besides the host genetics, it has the advantage of having a simpler bacterial community, when compared with mammals, and being relatively simple to produce axenic and gnotobiotic animals. *D. melanogaster* raised in axenic conditions have a delayed development, and are not viable under certain nutritional conditions, and bacteria can rescue these developmental problems [14-16]. Bacteria also affect the fly lifespan, gut homeostasis, interaction with pathogens, and behavior [17-23]. All of these phenotypes demonstrate the importance of bacteria to this host and the need to understand these interactions for a comprehensive view of *D. melanogaster* biology.

Despite the recognized importance of gut-associated bacteria to *D. melanogaster* what constitutes its gut microbiota is still an open question. Laboratory *D. melanogaster* is associated with few bacterial species, which belong mainly to *Acetobacter* and *Lactobacillus* genera [20,22,24-27]. This contrasts with data from flies sampled in their natural environment, which have a more diverse population of bacteria. In addition to *Acetobacter* and *Lactobacillus*, they are also enriched in bacteria from other families and genera [25,28]. Because *D. melanogaster* feeds on fermenting and rotten fruits containing many microbes, it is, however, difficult to understand which of the identified bacteria are colonizing the host gut and which are transiently passing with the food. Likewise, a similar problem is present in laboratory conditions, where flies live in a relatively closed environment. The bacteria found in their gut could simply correspond to food growing bacteria ingested by the flies. This hypothesis is supported by the fact that frequent transfer of adult flies to clean food vials strongly reduces their gut bacterial loads [20,27]. Consequently, the current working model is that the gut-associated bacteria in *D. melanogaster* are environmentally acquired and do not constitute *bona fide* gut symbionts.

Most functional studies in *D. melanogaster*, however, have been performed with bacterial isolates from lab stocks. The properties of bacterial isolates from wild-caught *D. melanogaster* could differ. Bacteria found in the gut of some other *Drosophila* species differ from the bacteria present in their food source, suggesting that they can be gut symbionts [29,30], and raising the possibility of these also existing in *D. melanogaster*. Moreover, a recent study compared the ability to colonize the gut of different *Lactobacillus plantarum* strains and found that one wild strain was able to colonize flies more frequently that strains isolated from laboratory flies [31]. Therefore, it is possible that natural populations of *D. melanogaster* have stable colonizing bacterial communities in their guts.

Here we analyzed bacterial isolates from the gut of wild-caught *D. melanogaster* and compared it to bacteria from lab stocks. Using a protocol that avoids re-infection of flies with bacteria growing on the food, we identified bacterial species that are stably associated with the gut of wild *D. melanogaster*. Moreover, these isolates can stably associate and proliferate in the gut of lab flies. We further analyze the specificity of these interactions and fitness advantage of stable associations. Our results lead to the identification of gut symbionts in *D. melanogaster* and demonstrate fitness advantages for both partners in an ecological context.

## Results

### Wild caught flies have stable gut-colonizing bacteria

In order to analyze the diversity and stability of gut bacteria in *Drosophila melanogaster* we used culture-dependent techniques. We plated single gut homogenates in agar plates of five different culture media (Luria Broth (LB), Mannitol, Brain Heart Infusion (BHI), MRS Broth (MRS), and Liver Broth Infusion (LBI)). This approach allowed us to determine absolute number of bacteria present in each gut and isolate bacteria for follow up experiments.

We started by analyzing levels of bacteria in the gut of flies from our standard laboratory stock *w^1118^* DrosDel isogenic strain (*w^1118^ iso*) [32,33]). We assessed these levels in young conventionally raised flies (Day 0) and after these flies were maintained singly for ten days either in the same vial or passed to a new vial twice a day (similarly to the protocol in [20]). The latter protocol was designed to decrease the probability of flies getting re-infected with their own bacteria or bacteria growing on fly food and, therefore, allowed us to test if there was a resident gut bacterial microbiota in this *D. melanogaster* lab stock (stability assay). In flies kept in the same vial for ten days, bacterial levels in the gut increased approximately 17-fold (Fig 1A and S1A Fig, linear mixed model fit (lmm), *p* < 0.001). In contrast, flies that were passed twice a day to a new vial, during these ten days, had an approximately 2,200-fold decrease in their gut bacterial levels (Fig 1B and S1A Fig, lmm, *p* < 0.001). A sharp decrease in bacterial loads was confirmed by quantitative PCR (qPCR), a culture independent method, using universal primers for the 16S rRNA gene (Fig 1C and S1B Fig, lmm, *p* < 0.001). These results show that bacterial levels in the gut of these flies are dependent on fly husbandry and suggest that these bacteria are transient, similarly to what was previously shown with a different laboratory stock [20]. Since these bacteria are associated with the lab stock and bacterial loads in the gut of these flies actually increase over time if they are kept in the same vials for ten days, we tested their growth on fly food (Fig 1D). We placed single flies per vial (Day 0), discarded them after 24 hours (Day 1), and kept the vials for a further nine days (Day 10). Bacterial levels on the surface of the fly food increased 7.6 x 10^8^ fold, from Day 1 to Day 10, clearly showing their capacity to grow on fly food (Fig 1D, lm, *p* < 0.001). Therefore, the bacteria associated with this lab stock grow on the fly food and are only transiently associated with the gut of adult flies.

**Fig 1.**
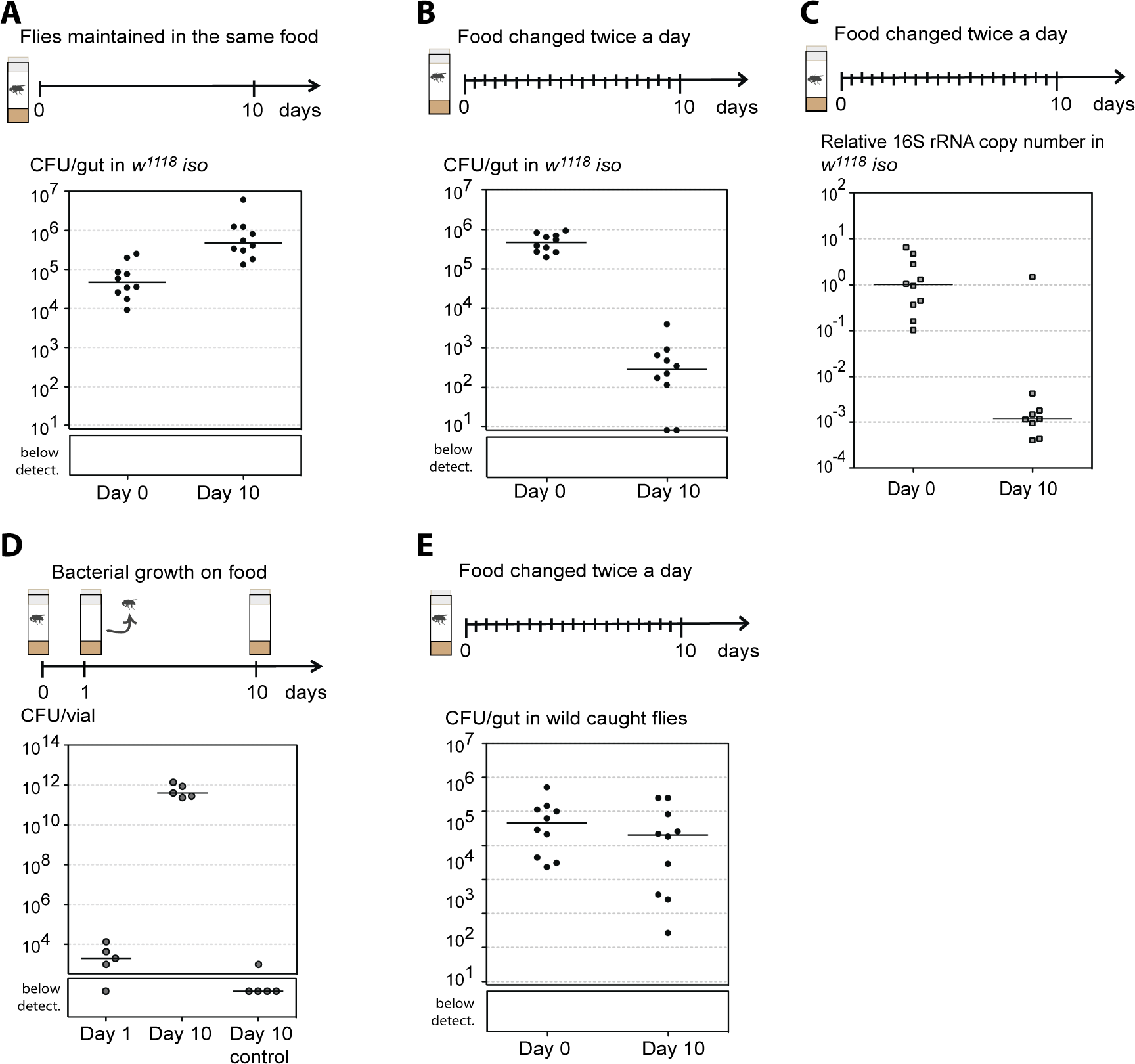
Wild-caught *D. melanogaster* have a stable gut microbiota. Single 3-6 days old *w^1118^ iso* males were kept in the same vial during ten days (A) or exposed to a stability protocol by being passed to new vials twice a day (B, C). (A, B) Ten individuals were analyzed at each day and total number of CFUs per gut determined by bacterial plating. Bacterial levels between day 0 and day 10 increase in (A) and decrease in (B) (lmm, *p* < 0.001 for both). (C) Relative amount of 16S rRNA bacterial gene was measured by quantitative-PCR in ten individual guts from each day, using the host gene *Rpl32* as a reference gene. Relative amount of 16S rRNA gene decreases between days (lmm, *p* < 0.001). (D) Single 3-6 days old *w^1118^ iso* males were placed in food vials for 24 hours and then discarded. Bacterial levels on the food were determined at this point (Day 1) and after incubating the vials for further nine days (Day 10). Bacterial levels were also assessed in control vials, not exposed to flies (Day 10 control). Five vials were analyzed for each condition and total number of CFUs per vial determined by bacterial plating. Bacterial levels increase between Day 1 and Day 10 (lm, *p* < 0.001). (E) Bacterial levels from wild-caught flies at the day of collection (Day 0) and after 10 days of the stability protocol (Day 10). Ten individuals were analyzed for each day and total number of CFUs per gut determined by plating. Bacterial levels on the flies significantly decrease with time (lmm, *p* = 0.004). (A-E) Each dot represents an individual gut or vial and lines represent medians. Statistical analyses were performed together with replicate experiments shown in S1 Fig.

We next asked if we could find stable bacteria in the gut of *D. melanogaster* collected from natural populations. We captured *D. melanogaster* from a population growing on fallen figs and quantified their gut bacterial levels at the time of collection (Day 0) and ten days after using the same stability assay designed to avoid re-infection (Day 10) (Fig 1E, S1C and S1D Fig). Although there is a statistical significant change in the bacterial levels in the gut with time (lmm, *p* = 0.004), the bacterial levels only decreased 4.8 fold in ten days. Moreover, at Day 10 wild flies maintained 2.9 x 10^4^ CFU per gut, while *w^1118^ iso* flies only had 100 CFU per gut. Also, even after 20 days of this protocol wild flies still maintained approximately 6.1 x 10^3^ CFU per gut (S1D Fig), showing a long-term stability of their microbiota. These results show that wild flies carry bacteria that are stably associated with their gut.

In order to identify and isolate the bacteria that can stably interact with the gut of *D. melanogaster*, we analysed the bacterial composition of the cultured gut extracts of *w^1118^ iso* and wild flies represented in Fig 1B and 1E. For each fly gut homogenate, in each of the five media, we distinguished colonies by morphology, determined CFUs per gut of each morphological type, and isolated two colonies of each morphological type. For each isolate we sequenced by Sanger a fragment of the 16S rRNA gene, which included the V2 to V4 hypervariable regions. After sequencing we classified morphological types into operational taxonomic units (OTUs), based on Greengenes alignment tool and database [34], and determined the number of colony forming units (CFUs) of each OTU in each fly gut (Fig 2). In general we could assign each morphological type to one OTU. However, in samples from wild flies we could not distinguish by morphology the colonies of different *Lactobacillus* species, different Acectobacteraceae (*Acetobacter* and *Gluconobacter* species), and several genera of Enterobacteriaceae. We therefore calculated CFUs per fly for each of these groups of bacteria and not individual OTUs (Fig 2). The frequencies of the different OTUs belonging to these groups, in the different conditions, are shown in Fig 3B, 3D, 3F, 3H and S3 Fig.

**Fig 2.**
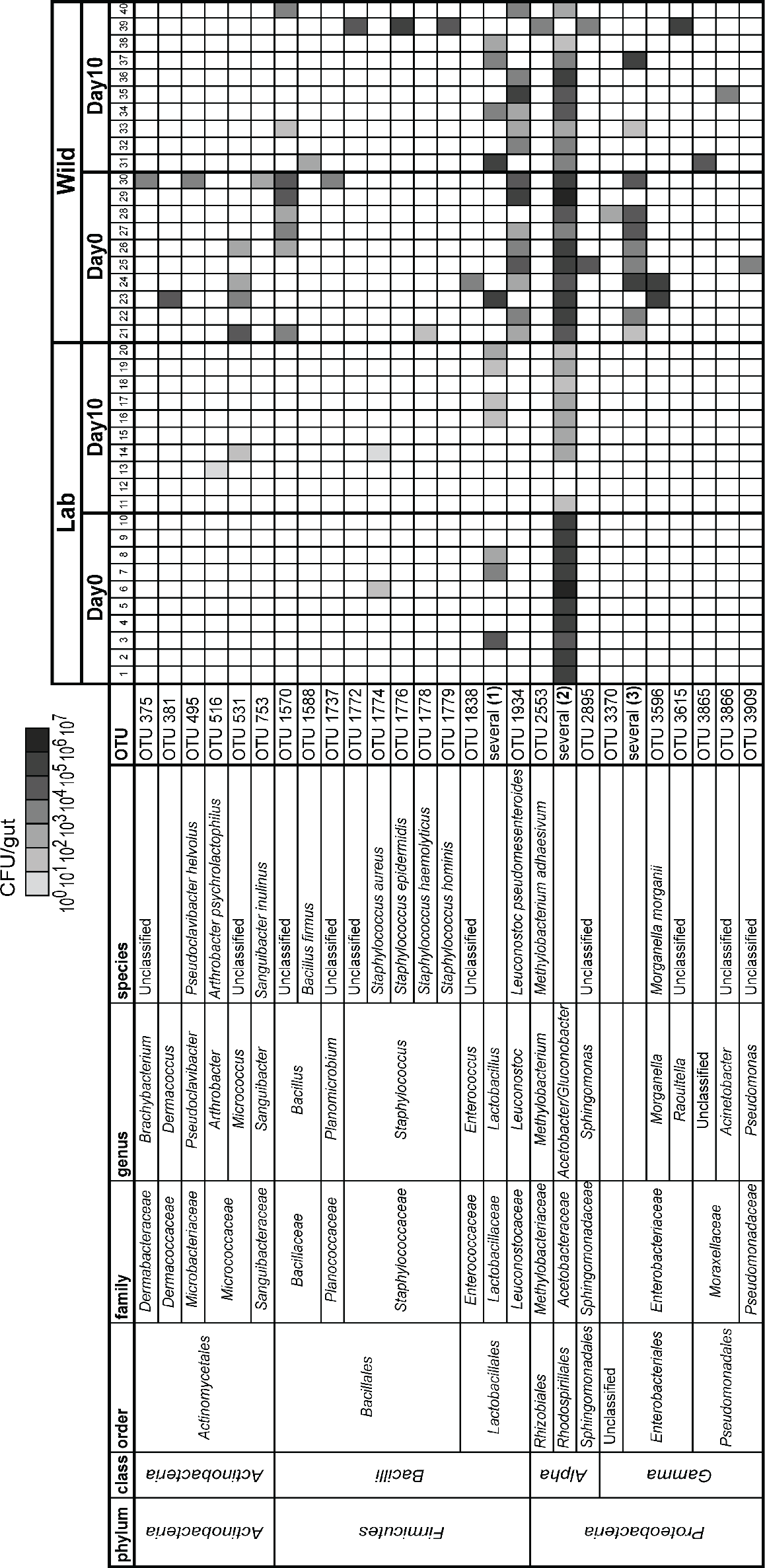
Higher diversity of gut bacterial communities in wild-caught *D. melanogaster*. Bacterial OTUs present in the gut of laboratory (1-20) and wild caught (21-40) flies before (Day 0) and after being exposed to the stability protocol (Day 10). Gut homogenates from flies represented in Fig 1B and E were plated in different culture media and representative colonies of each morphological type were sequenced. Each column represents one individual gut. Bacterial levels are represented on a grey-scale from 10^0^ to 10^7^ CFUs per gut. Colonies of different *Lactobacillus, Acetobacteriaceae* or *Enterobactereaceae* were not possible to distinguish by morphological type and are grouped together. The presence of *Lactobacillus* species and *Leuconostoc pseudomesenteroides* in wild-caught flies is not independent (Pearson’s Chi-squared test, *p* = 0.014). Frequencies of the different OTUs in these groups are represented on Fig 3B, D, F, H and S3B Fig.

**Fig 3.**
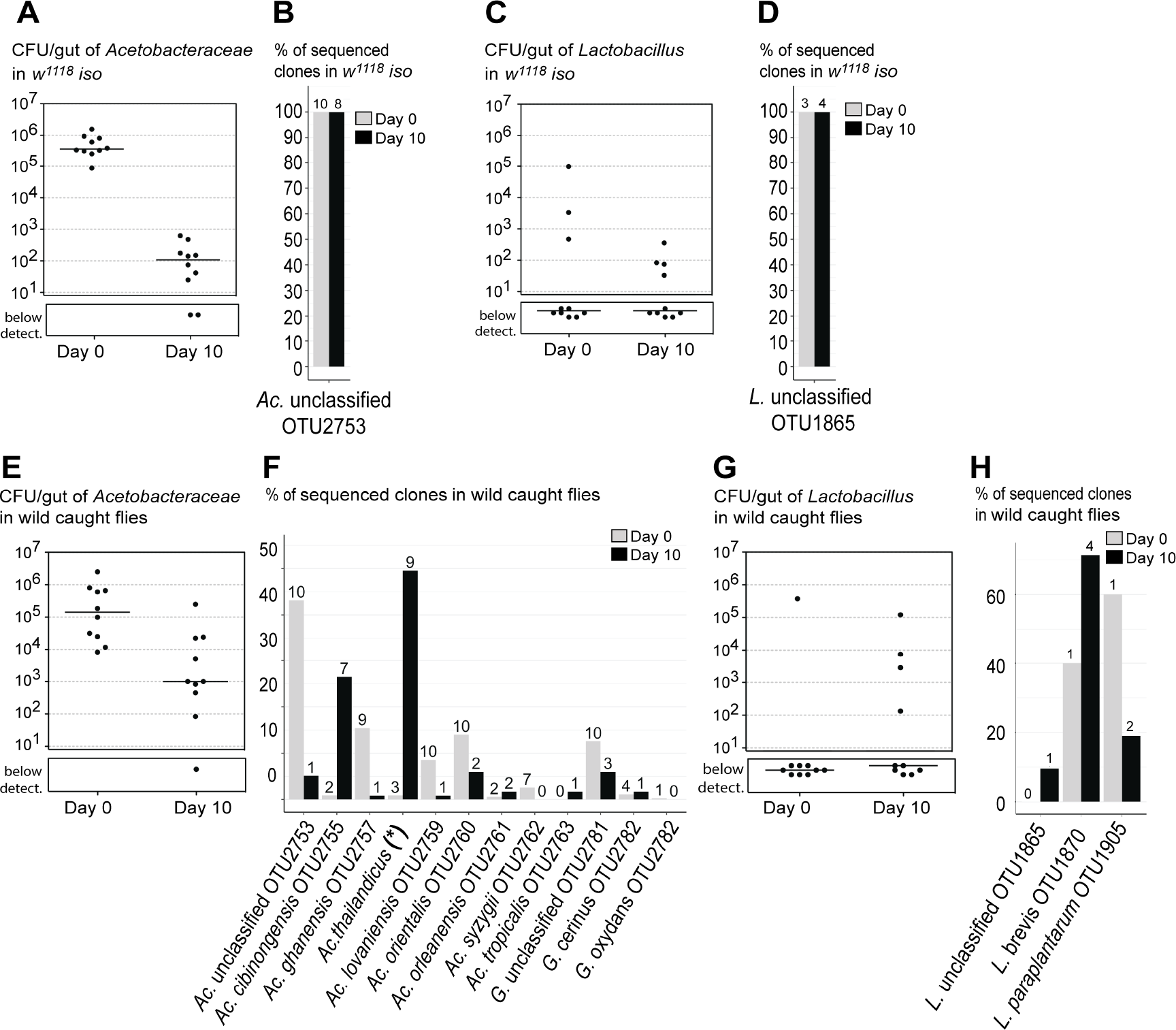
Wild-caught flies maintain in the gut particular *Acetobacter* species. Total levels of *Acetobactereaceae* (A, E) and *Lactobacillus* (C, G) in laboratory *w^1118^ iso* (A, C) and in wild-caught flies (E, G) before (Day 0) and after 10 days of the stability protocol (Day 10). Each dot represents one individual gut and lines represent medians. Levels of *Acetobactereaceae* decrease between days in both types of flies (lm, *p* <= 0.002 for both). Changes in levels of *Lactobacillus* are not significant in both (lm, *p* >= 0.302). Frequencies of sequenced colonies of *Acetobactereaceae* (B, F) and *Lactobacillus* (D, H) in *w^1118^ iso* (B, D) and in wild-caught flies (F, H). *Ac.- Acetobacter*, *G. - Gluconobacter* and *L. - Lactobacillus*. Numbers on the top of the bars correspond to the number of 10 flies carrying each OTU, from a total of flies (B, D, F, H). * *Ac. thailandicus* was initially identified as *Ac. indonesiensis* OTU2758 based on partial sequence of 16S rRNA gene.

Laboratory flies presented very low diversity in their gut bacterial community, as previously reported [25,26,28]. From each gut of laboratory flies we could isolate one to two different OTUs at Day 0, and zero to three different OTUs at Day 10. In total, we isolated from these flies three and five different OTUs at Day 0 and Day 10, respectively (S2 Fig). Also, the accumulation curves indicate that we sampled most of the diversity present in laboratory flies possible with our approach (S2 Fig). Laboratory flies were mainly found associated with two OTUs, *Acetobacter* OTU2753 and *Lactobacillus* OTU1865 (Fig 2, Fig 3A-3D). On Day 0, all the flies were associated with high levels of *Acetobacter* OTU2753 (Fig 3A, 3B), while *Lactobacillus* OTU1865 was only present in some individuals (Fig 3C, 3D). After 10 days of the stability assay, *Acetobacter* levels decrease (lm, *p* = 0.001), while *Lactobacillus* levels are not significantly different *(p =*0.635) (Fig 3A, 3C). Importantly, when we analyzed the bacterial species that were capable of growing on fly food in Fig 1D, we found these two same OTUs, with *Acetobacter* OTU2753 being the most abundant. Altogether, these results show that this *D. melanogaster* laboratory stock has very low bacterial diversity and is mainly associated with transient bacteria able to grow on fly food.

In contrast, wild caught flies were associated with a higher diversity of bacterial species (Fig 2, Fig 3F, 3H, and S3B Fig). From each gut of wild flies we isolated nine to 16 different OTUs at Day 0, and three to 14 different OTUs at Day 10. In total, we isolated 35 and 31 different OTUs at Day 0 and Day 10, respectively (S2 Fig). Moreover, it seems that we are not close to saturation with these samples and that further sampling would allow the identification of more OTUs associated with the gut of *D. melanogaster* from this wild population (S2 Fig).

The individual characterization of bacterial species present in each gut allowed us to discriminate between OTUs that were only present in one or few individuals, albeit at higher levels, and OTUs associated with most individuals. At the day of collection (Day 0) 50% or more of the flies had in their gut *Bacillus* OTU1570, *Leuconostoc pseudomesenteroides* OTU1934, *Acetobacter* OTU2753, *Ac. ghanensis* OTU2757, *Ac. lovaniensis* OTU2759, *Ac. orientalis* OTU2760, *Ac. syzygii* OTU2762, *Gluconobacter* OTU2781, *Enterobactereaceae* OTU3529, *Tatumella* OTU3635 and *Kluyvera ascorbata* OTU3643 (Fig 2, Fig 3F and S3B Fig). Ten days after the stability assay only a few bacteria remained associated with the gut of most individuals. One of these bacteria was *L. pseudomesenteroides*, which was present in six out of ten flies and did not show a significant reduction in levels between Day 0 and Day 10 (Fig 2, S4 Fig lm, *p =* 0.372). Bacteria from the *Acetobacteraceae* family also remained associated with the gut of most wild flies, at an estimated 1.3 x 10^3^ CFU per gut at Day 10, despite a significant reduction of approximately 100-fold in their levels between Day 0 and Day 10 (lm, *p* = 0.002) (Fig 2, Fig 3E). However, the frequencies of different OTUs of *Acetobacteraceae* changed significantly between Day 0 and Day 10 (Fig 3F, Pearson’s Chi-square with Monte Carlo simulation, *p* < 0.001). At Day 10, all the OTUs that were dominant at Day 0 become present at lower frequencies and *Acetobacter cibinongensis* OTU2755 and *Acetobacter indonesiensis* OTU2758 (later re-identified as *Acetobacter thailandicus*, see below) became the dominant bacteria (Fig 3F). These two bacteria were present in at least seven and nine individuals out of ten, respectively, and together represented 76% of the sequenced colonies.

We isolated clones of these gut bacteria to further characterize them. Analysis of the full 16S rRNA gene sequence (S1 Text) confirmed the identity of the wild isolates *Leuconostoc pseudomesentoroides* OTU1934 and *Acetobacter cibinongensis* OTU2755, based on Greengenes [34]. We also confirmed the identity of the transient laboratory isolate as *Acetobacter* OTU2753. However, the analysis of the full 16S rRNA gene from the previously identified *Ac. indonesiensis* OTU2758 isolate matched several different *Acetobacter* OTUs with 98% identity. Therefore, we used BLAST to analyze the full sequence against the NCBI 16S ribosomal RNA sequences (Bacteria and Archaea) database [35]. *Ac. thailandicus* 16S rRNA gene was the best hit and was 99% identical to the sequence of this isolate [36].

Overall, this analysis identified three species that seem to be stably associated with the gut of wild flies in this population: *L. pseudomesenteroides, Ac. cibinongensis* and *Ac. thailandicus*.

#### *Acetobacter thailandicus* and *Ac. cibinongensis* proliferate in the gut of *D. melanogaster* and are stably associated with it

To study the interaction of these bacteria with *D. melanogaster* we generated stocks of *w^1118^ iso* flies monoassociated with each of these bacteria and we tested their persistence using the stability assay. In agreement with our previous observations, the laboratory isolate of *Ac*. OTU2753 did not persist in the gut and disappeared from the majority of the flies (lmm, *p* < 0.001) (S5A and S5E Fig). On the other hand, the wild isolates of *Ac. cibinongensis, Ac. thailandicus* and *L. pseudomesenteroides* persisted in the gut of flies until Day 10, showing a more stable association with the host (S5B-S5D and S5F-S5H Fig). *L. pseudomesenteroides* levels did not significantly change with treatment *(p =* 0.96) and, although *Ac. cibinongensis* and *Ac. thailandicus* levels significantly decreased in the ten days (*p* < 0.001 for both), both remained in the gut at approximately 100 and 3,800 CFUs, respectively.

To better assess the bacterial dynamics within the gut, we developed a more strict protocol to avoid re-infection. We maintained single flies in cages with a larger food surface (356.7 cm^2^ compared with 3.8 cm^2^ in vials), which was changed daily (Fig 4A). We assessed gut bacterial levels at the beginning of the experiment and after one, two, five and ten days of this treatment. In accordance with previous data, *Ac*. OTU2753 levels rapidly decreased and most flies had no detectable bacteria in their gut after five days of treatment (Fig 4B). *Ac. cibinongensis* and *Ac. thailandicus* also presented an initial decrease in bacterial levels in the gut, but these seemed to stabilize after two days of treatment, confirming their stability in the gut (Fig 4C and D). However, and contrary to what was observed in vials, *L. pseudomesenteroides* was not stable when the protocol was performed in cages (Fig 4E). After two days, approximately 50% of flies lost *L. pseudomesenteroides* from their gut. An independent replicate with data from only Day and Day showed similar results for all bacteria (S5E-S5H Fig).

**Fig 4.**
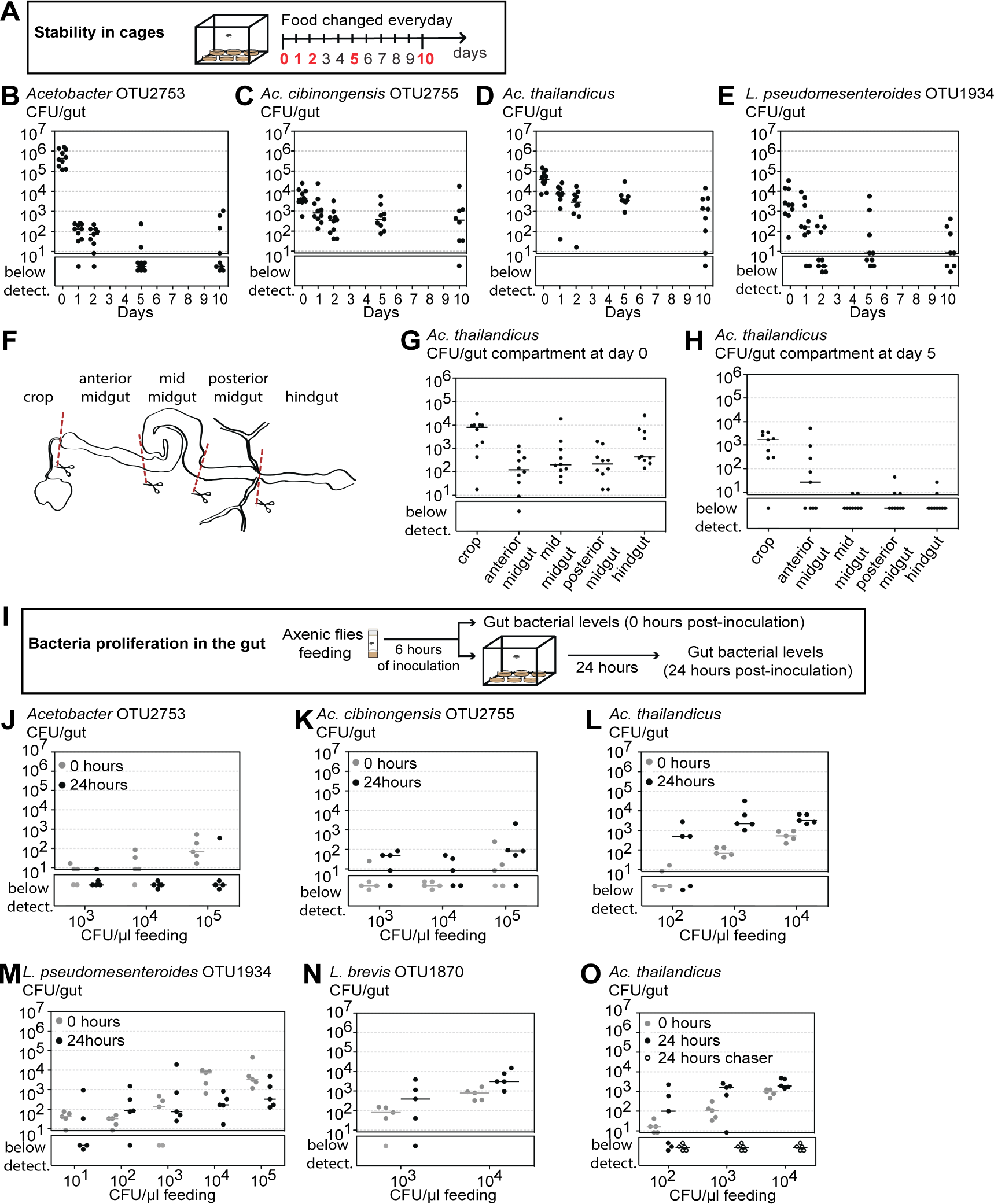
*Ac. thailandicus* and *Ac. cibinongensis* proliferate and stably colonize the gut of *D. melanogaster*. (A-E) Stability of different bacteria in monoassociation. Single 3-6 days old *w^1118^ iso* males from monoassociated stocks with *Ac*. OTU2753 (B), *Ac. cibinongensis* (C), *Ac. thailandicus* (D) or *L. pseudomesenteroides* (E) were exposed to the stability protocol in cages, as shown in the scheme (A). Number of CFUs in individual guts was assessed by plating at days 0, 1, 2, 5 and 10 of the protocol. Stability of different bacteria was analyzed by fitting the data to an exponential decay model represented in S5I Fig. (F-H) Distribution of *Ac. thailandicus* in the gut. Scheme of gut regions analysed (F). Number of CFUs in each gut compartment from *w^1118^ iso* males monoassociated with *Ac. thailandicus* before (G) and after (H) five days of the stability protocol. (I-O) Proliferation of different bacteria in the gut of *D. melanogaster*. 3-6 days old axenic *w^1118^ iso* males were inoculated for 6 hours with different concentrations of *Ac*. OTU2753 (J), *Ac. cibinongensis* (K), *Ac. thailandicus* (L, O), *L. pseudomesenteroides* (M), and *L. brevis* (N). Bacterial levels were assessed 0 and 24 hours post-inoculation. During this period males were singly placed in cages as shown in the scheme (I). In (O) axenic chaser males were placed in cages together with males inoculated with *Ac. thailandicus*. At 24 hours bacterial levels were assessed for both males. Bacterial levels between 0 and 24 hours decrease in flies inoculated with *Ac*. OTU2753 (lmm, *p* < 0.001), increase in flies inoculated with *Ac. cibinongensis, Ac. thailandicus*, and *L. brevis* (*p* = 0.024, *p* < 0.001, and *p* = 0.046, respectively) and do not significantly change in flies inoculated with *L. pseudomesenteroides* (*p* = 0.158). Ten (B-E and G-H) or five (J-O) individuals were analyzed for each condition, per replicate, and total number of CFUs per gut determined by plating. Each dot represents one gut or one gut fragment and lines represent medians. Statistical analyses were performed together with replicate experiments shown in S5 and S6 Fig.

We compared the dynamics of the gut levels of the four bacteria by fitting the data of Fig 4B-4E to an exponential decay model (S5I Fig). This model estimates the exponential decay rate which corresponds to the rate of bacterial loss from the gut and an asymptote that corresponds to the levels at which the bacteria tend to stabilize after this loss. The simplest model that explains the data has the same estimate for the exponential decay rate for all the bacteria. There are, however, significant differences between the asymptotes of all the bacteria (Contrasts between nonlinear least-square estimates, *p* < 0.014), except between *Ac*. OTU2753 and *L. pseudomesenteroides (p =* 0.395). Overall, an interpretation of this fit is that in all cases most of the bacterial population is in an unstable compartment, at the beginning of the experiment, from where they tend to disappear with similar dynamics. However, *Ac. cibinongensis* and *Ac. thailandicus* are also present in a stable compartment, at levels that correspond to the calculated asymptotes (approximately 300 and 1,300 CFU per gut, respectively).

In order to identify in which gut region bacteria could be stably associated with the host, we analyzed *Ac. thailandicus* levels present in different gut regions before (Day 0) and after 5 days of the stability protocol in cages (Day 5) (Fig 4F). At Day 0, *Ac. thailandicus* was distributed along the gut, being present at lower levels in the midgut, compared with crop and hindgut (Fig 4G and S5J Fig). After 5 days, bacteria were found in two anterior gut sections, one comprising the crop and the other comprising the anterior midgut and the proventriculus (Fig 4H and S5K Fig). Therefore, the niche for the stable population of *Ac. thailandicus* is the anterior part of the gut.

We next asked if these bacteria had the capacity to proliferate in the gut of *D. melanogaster*, since stability in the gut could be achieved through other mechanisms (e.g. bacteria could be simply attaching to the gut and avoiding elimination). Thus, we developed a protocol to analyze proliferation based on giving a small inoculum of bacteria and test if bacterial loads increase over 24h. We raised flies in axenic conditions and exposed 3-6 days old males to different doses of bacteria. After 6 hours of feeding on the bacteria inoculum, flies were either collected to dissect and assess bacterial levels in the gut (0h) or placed singly in cages, as described above, and collected 24 hours later (Fig 4I). In this assay, *Ac.* OTU2573 did not colonize the gut of adult flies and at the higher inoculum titers the levels decreased between 0h and 24h (lmm, *p* < 0.001) (Fig 4J, S6A, S6E Fig), indicating that these bacteria cannot proliferate in the gut of *D. melanogaster*. On the other hand, the levels of *Ac. cibinongensis* and *Ac. thailandicus* increased in 24h (*p* = 0.024 and *p* < 0.001, respectively) (Fig 4K, 4L, S6B, S6C, S6F, S6G Fig), showing that these bacteria can proliferate in the gut of *D. melanogaster. Ac. thailandicus* proliferate more and reached higher levels than *Ac. cibinongensis* (*p* = 0.019). Interestingly, in flies exposed to *Ac. thailandicus* inoculums superior to 10^2^ CFU/µl, these bacteria reach between 600 and 1,900 CFU per gut (Fig 4L, S6C and S6G Fig). These levels are similar to the stable compartment population size estimated above (1,300 CFU per gut), indicating that *Ac. thailandicus* can rapidly colonize a fly.

*L. pseudomesenteroides* levels did not significantly increase or decrease over 24h (lmm, *p =* 0.158) (Fig 4M and S6D Fig). At inoculums superior to 10^2^ CFU/µl, *L. pseudomesenteroides* levels at 24h are between 150 and 550 CFU per gut. These results fail to show proliferation of *L. pseudomesenteroides* but indicate that this bacterium is not eliminated at the same rate as the unstable *Ac*. OTU2753.

Since *Lactobacillus* species are commonly found associated with *D. melanogaster* and shown to impact its physiology [25,26,28,37-39], we also tested isolates of *Lactobacillus paraplantarum* OTU1905 and *Lactobacillus brevis* OTU1870 in this assay (Fig 4N, S6H-S6J Fig). These *Lactobacillus* were isolated from the gut of wild flies at day 0 of the stability assay (Fig 3G and 3H) and the isolates identity confirmed by sequencing the full 16S rRNA gene (S1 Table). *L. paraplantarum* levels do not change over 24h (lmm, *p* = 0.65) and can be sustained at 200 to 800 CFU per gut (similarly to *L. pseudomesenteroides)* (S6H and S6I Fig). On the other hand, the levels of *L. brevis* increase in 24h *(p =* 0.046), showing that this bacterium proliferates in the gut of *D. melanogaster* (Fig 4N and S6J Fig).

Overall, these assays show that *Ac. cibinongensis, Ac. thailandicus*, and *L. brevis* isolates proliferate in the gut of *D. melanogaster*. On the contrary, the transient *Ac.* OTU2753 cannot proliferate and is rapidly lost. *L. pseudomesenteroides* and *L. paraplatarum* have an intermediate phenotype where proliferation is not shown but the bacteria can sustain themselves in the gut over a period of 24h after oral inoculation.

As all these *Acetobacter* species were able to grow on fly food (S7 Fig), it was still possible that the increase in the levels of *Ac. thailandicus* in the proliferation assay (Fig 4L, S6C Fig) was due to a very fast growth on the fly food and re-acquirement by feeding. To test this possibility we placed axenic (chaser) flies in cages simultaneously with the flies that had fed on *Ac. thailandicus*, at time 0h of the experiment. At 24 hours none of the axenic chaser flies had bacteria in their gut (Fig 4O and S6G Fig). This demonstrates that the levels measured in the inoculated fly were due to proliferation in the gut and not due to bacteria acquired from the food.

#### *Ac. thailandicus* gut proliferation is species specific

To test if proliferation of *Ac. thailandicus* in the gut is host specific we compared its proliferation in *D. melanogaster* and *Drosophila simulans*. These two species share the same habitat, feed on the same material and are frequently captured together [40]. We used a proliferation protocol similar to the one described above (see figure legend, S8A, S8B Fig) to test three different genetic backgrounds of each host species. These included one isofemale line of each species that were collected simultaneously, from the same place as the initial collection of wild *D. melanogaster*. There is a significant difference in the colonization by *Ac. thailandicus* in these two host species (Fig 5, S8C-S8E Fig, lmm, *p* < 0.001), with the levels increasing over 24h in *D. melanogaster* but decreasing in *D. simulans*. These results suggest that *D. melanogaster* and *Ac. thailandicus* interaction is host specific. Interestingly, although *Ac. thailandicus* colonizes all strains of *D. melanogaster* tested (Fig5, S8C-S8E Fig), there is variation in the growth at 24h, indicating modulation of this process by the host genotype (lmm, *p* = 0.002).

**Fig 5.**
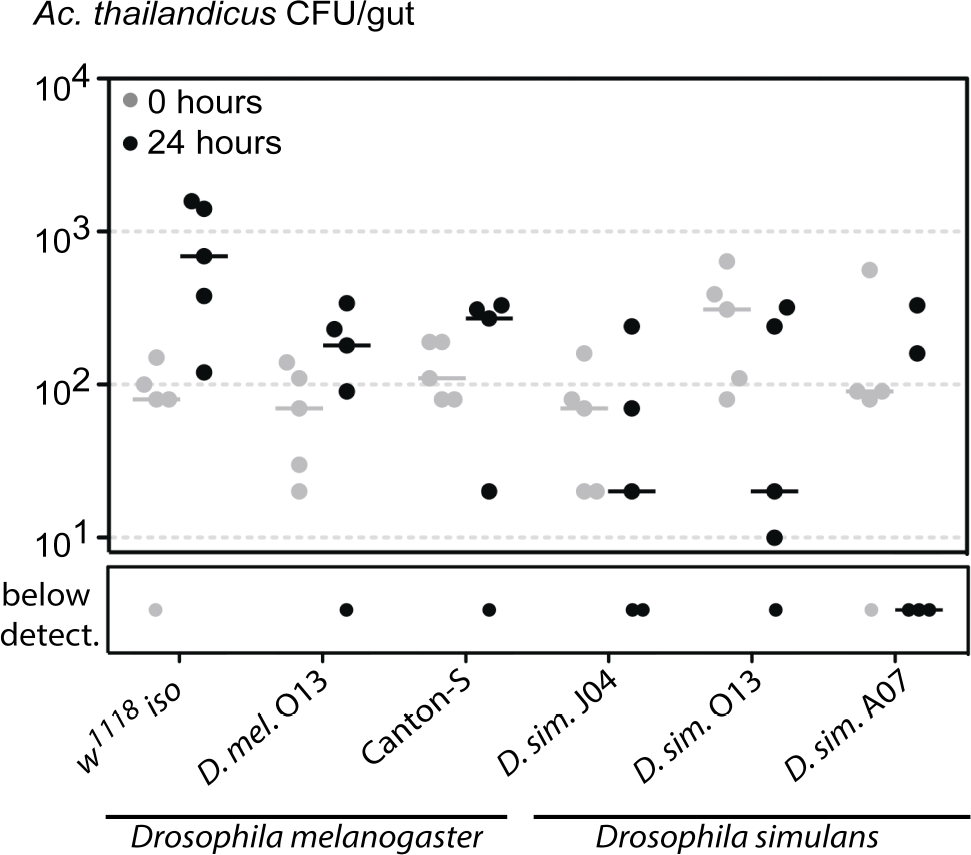
*Ac. thailandicus* proliferates specifically in *D. melanogaster* but not in *D. simulans*. Axenic 3-6 days old *D. melanogaster* or *D. simulans* males were inoculated for 6 hours with *Ac. thailandicus* (10^4^ CFU/µl). Bacterial levels were assessed 0 and 24 hours post-inoculation. During this period males were singly placed in bottles. Three different genetic backgrounds for *D. melanogaster (w^1118^ iso*, *D. mel*. O13 and Canton-S) and for *D. simulans (D. sim*. J04, *D. sim*. O13 and *D. sim*. A07) were tested. Bacterial levels in the gut increase in *D. melanogaster* and decrease in *D. simulans* (lmm, *p* < 0.001). Five individuals were analyzed for each condition and total number of CFUs per gut determined by plating. Each dot represents one gut and the lines represent medians. Statistical analysis was performed together with replicate experiments shown in S8C-E.

#### *Ac. thailandicus* stable association with *D. melanogaster* is mutually beneficial

Symbiotic associations can range from pathogenic to mutualistic. As *Acetobacter* species have been previously described as beneficial to *D. melanogaster* [16] we tested if the stable association between *D. melanogaster* with *Ac. thailandicus* could be advantageous for both. We started to test this hypothesis by comparing fitness parameters of flies monoassociated with *Ac. thailandicus, Ac*. OTU2753 and axenic flies by measuring time to pupariation and adulthood and total number of its progeny. Both *Ac. thailandicus* and *Ac*. OTU2753 monoassociated stocks had a much higher fertility than axenic flies and there was no significant difference between them (S9A, S9B Fig, lm, *p* < 0.001 for the comparisons of each *Acetobacter* monoassociation with axenic flies, in number of pupae or adults, *p* > 0.968 for the comparisons between *Acetobacter* monoassociated stocks). Flies monoassociated with either *Acetobacter* also developed until pupariation or adulthood approximately 3 days faster than axenic flies (S9C, S9D Fig, lm, *p* < 0.001 for each *Acetobacter* monoassociation comparison with axenic flies). Flies monoassociated with *Acetobacter* OTU2753 developed slightly faster to pupae (0.38 days) and adults (0.57 days) (*p* < 0.001 for each comparison). These results show that in this setup the association with either *Acetobacter* is clearly advantageous when comparing with axenic conditions and that the stable *Ac. thailandicus* does not provide a greater benefit than the lab isolate *Ac*. OTU2753.

However, the advantage of a stable association may not be revealed by directly studying monoassociated *D. melanogaster* stocks. In these conditions the bacteria are continuously associated with *D. melanogaster*, even if it only present in the food or transiting through the gut. But in the wild *D. melanogaster* adults freely move in space and can explore a continuously changing environment, a situation in which a stable association could be important. Therefore, we established a protocol to test the fitness benefits of the stable interaction in a scenario that simulates this changing environment. The protocol is similar to the proliferation protocol outlined above. After six hours of feeding on an inoculum of bacteria, one female and two males were placed per cage and maintained there for ten days, with food being changed daily (Fig 6A). After ten days of this protocol males exposed to *Ac. thailandicus* have a median of 6,800 CFU per gut (Fig 6B and S10A Fig), showing that colonization can be sustained for a long time. In females, *Ac. thailandicus* grows in the gut between the beginning of the experiment and ten days in the cage (Wilcoxon rank sum test, *p* < 0.001) and reaches a median of 17,500 CFU per gut. These results show that *Ac. thailandicus* also colonizes and proliferates in female *D. melanogaster*. On the other hand, *Ac*. OTU2753 levels decrease between the beginning of the experiment and day ten in females *(p* = 0.048) and they have a median of 0 CFU per gut at day ten in both sexes, confirming that flies are not colonized by these bacteria (Fig 6B and S10A Fig).

**Fig 6.**
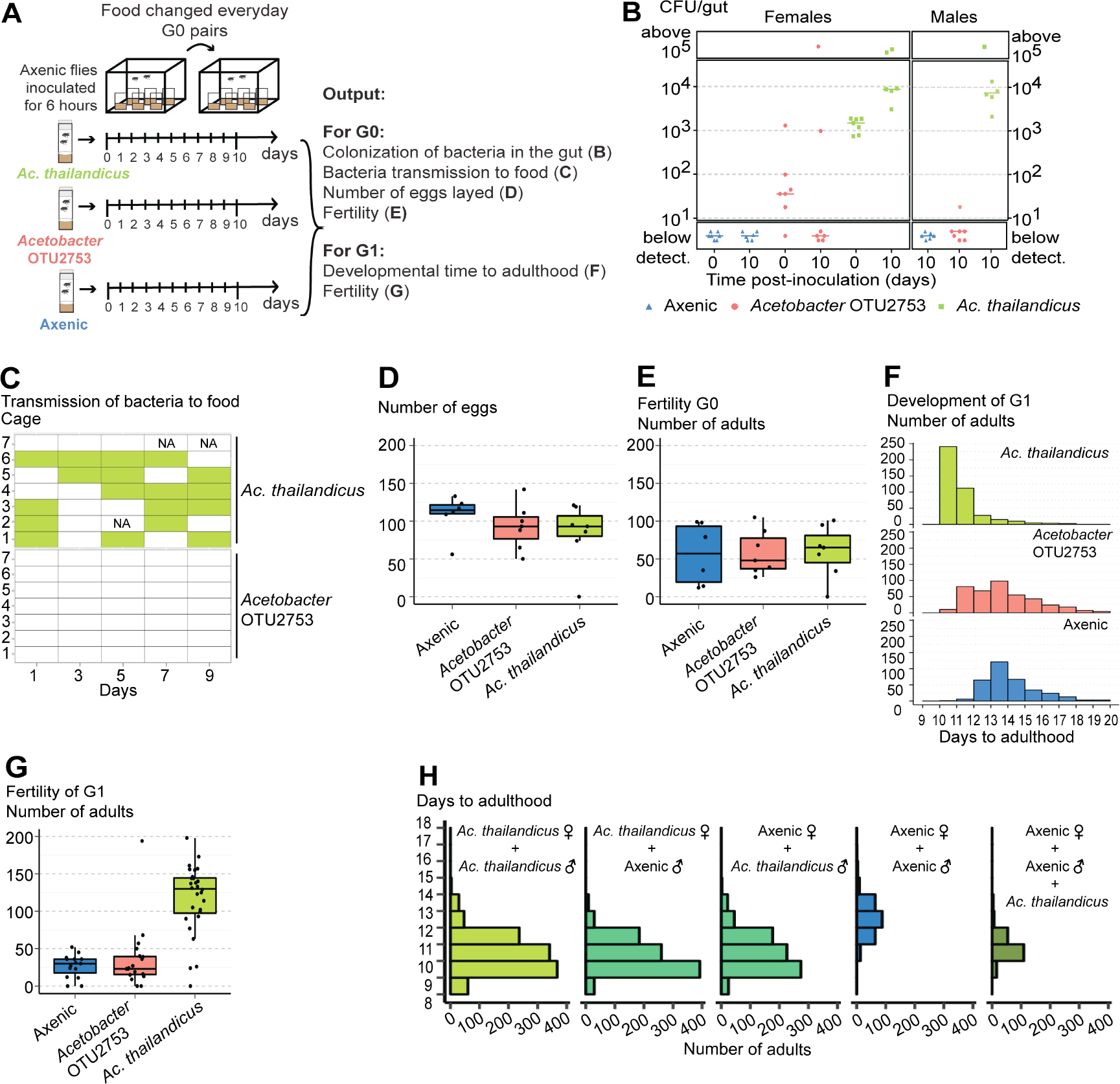
*Ac. thailandicus* stable association with *D. melanogaster* is mutualistic. (A) Axenic 1-3 days old *w^1118^ iso* males and females (G0) were in contact with an inoculum of 10^5^ CFU/µl of *Ac*. OTU2753, *Ac. thailandicus*, or sterile Mannitol (Axenic), for 6 hours. Two males and one female were placed per cage, with 6-7 cages for each condition, during 10 days with daily changed food. This experimental setup corresponds to data shown in panels B-G. (B) Bacterial levels in single guts of females at time 0 (0 days) and 10 days post-inoculation and in males 10 days post-inoculation, analyzed by plating. Bacterial levels between the two time-points increased in females inoculated with *Ac. thailandicus* and decreased in females inoculated with *Ac*. OTU2753 (Mann-Whitney test, *p* < 0.001 and *p* = 0.048 respectively). (C) Presence of bacteria on the food collected from cages at days 1, 3, 5, 7 and 9 of the protocol, analyzed by plating. Filled rectangles represent presence of bacteria. NA stands for samples that were not analyzed. *Ac. thailandicus* is transmitted to the food with higher frequency than *Ac*. OTU2753 (glm-binomial, *p* < 0.001). (D-G) Effect of bacterial association on the fitness of *D. melanogaster*. Total number of eggs laid by flies inoculated, or not, with different *Acetobacter* (D) and total number of adults that emerged from these eggs (E). Total number of eggs or adults is not different between conditions (lmm, *p* > 0.484 for all comparisons). (F) Developmental time to adulthood of the progeny (G1) of flies inoculated or not with different *Acetobacter*. Developmental time to adulthood is faster in progeny from flies inoculated with *Ac. thailandicus* than in the other two conditions and in progeny from flies inoculated with *Ac*. OTU2753 compared to progeny from axenic flies (lmm, *p* < 0.001, for these comparisons). (G) Fertility of G1 was assessed by placing two males and one female of G1 per vial, flipping them every other day for 10 days, and analyzing total number of emerged adults. Fifteen or more couples were made per condition. Fertility is higher in progeny from flies inoculated with *Ac. thailandicus* compared with the other two conditions (lmm, *p* < 0.001, for both comparisons) and not different in the comparison between the progeny of flies inoculated with *Ac*. OTU2753 or axenic (*p* = 0.592). (H) One male and one female 1-2 days old *w^1118^ iso*, either axenic or monoassociated with *Ac. thailandicus*, were placed in vials and flipped every other day for 10 days. To one set of vials with axenic parents *Ac. thailandicus* was added on the eggs after passing the parents. Developmental time to adulthood of the progeny was assessed. Ten couples were made per condition. There are no differences on developmental time to adulthood if either or both parents are monoassociated with *Ac. thailandicus* (lmm, *p* > 0.412 for all these comparisons). Progeny from couples where either or both parents are monoassociated and progeny from axenic flies where *Ac. thailandicus* culture is added on the eggs develop faster than progeny from axenic flies (lmm, *p* < 0.001, for all these comparisons). (B) Each dot represents one gut and lines represent medians. (D, E and G) Each dot represents the total progeny of one female. All statistical analyses were done together with replicate experiments shown in S10 and S11.

As a measure of the fitness benefit for the bacteria, in being stably associated with *D. melanogaster*, we tested if they could be transmitted to the food. We analyzed bacterial transmission by flies during the experiment, at days one, three, five, seven and nine. Flies associated with *Ac. thailandicus* transmitted bacteria to the food with a much higher frequency than flies associated with *Acetobacter* OTU2753, where transmission occurred only once (Fig6C, S10B Fig, generalized linear model with binomial distribution (glm-binomial), *p* < 0.001). Moreover, the probability of transmission of *Ac. thailandicus* to the food was independent of the day of the experiment (anova on glm-binomial models, *p* = 0.811). These results show that upon gut colonization *Ac. thailandicus* can be continuously transmitted by *D. melanogaster*. This may be advantageous to the bacteria and mediate their dispersal in the environment.

To compare the effect of this association on the host fitness, we started by analyzing the fertility of the flies in terms of number of eggs laid and adult progeny, during the experiment. The number of eggs or adult progeny were not significantly different between axenic flies and flies exposed to either bacteria (Fig 6D, 6E, S10C, S10D Fig, lmm, *p* > 0.484 for all comparisons). However, the time that these embryos took to reach adulthood was different. Progeny from flies colonized by *Ac. thailandicus* developed two or three days faster than progeny from flies previously exposed to *Ac*. OTU2753 or axenic flies, respectively (Fig 6F, S10E, lmm, *p* < 0.001 for both comparisons). However, the progeny of flies exposed to *Ac*. OTU2753 developed only 0.6 days faster than axenic flies (*p* < 0.001). Moreover, the fertility of this progeny was strongly influenced by the interaction of their parents with bacteria. The progeny from flies previously colonized by *Ac. thailandicus* had a much higher fertility than the progeny from flies previously exposed to *Ac*. OTU2753 or axenic flies (Fig 6G, S10F Fig, lmm, *p* < 0.001 for both comparisons), while there was no difference between the progeny of flies exposed to *Ac*. OTU2753 or axenic flies (*p* = 0.592). These data show that the interaction of adult flies with stable bacteria does not affect their fertility but has a strong influence on the development and fertility of its progeny.

This trans-generational effect could be due to an effect of the stable *Ac. thailandicus* gut population on the parents, and a subsequent indirect effect on the progeny, or through the transmission of the bacteria to the next generation and its effect during larval development. We tested if the developmental time of the progeny was dependent on the bacterial association with either parent by analysing the four possible couple combinations of flies raised axenically or monoassociated with *Ac. thailandicus* (Fig 6H, S11 Fig). There is no difference in developmental time to pupariation or adulthood if either or both parents are from the monoassociated stock (lmm, *p* > 0.412 for all these comparisons). The progeny of these three crosses develop, on average, 2.7 to 2.8 days faster than the progeny of crosses with both parents axenic (*p* < 0.001 for all comparison). These results show that the trans-generational effect on developmental time is not specifically associated with the mother or the father. Also, adding *Ac. thailandicus* to the progeny of axenic flies rescues the developmental delay. When bacteria are added these flies develop approximately two days faster (*p* < 0.001). This is not a full rescue since axenic eggs plus *Ac. thailandicus* still develop, on average, 0.5 to 0.8 days slower than flies with either or both parents from monoassociated stocks (*p* < 0.001 for all comparisons). This may be explained by the fact that the bacteria are only added when the parents are removed from the vial, after two days of egg laying. These data is compatible with a scenario where flies associated with *Ac. thailandicus*, either male or female, can transmit the bacteria to the next generation, which then plays an important role in its development. In agreement with this hypothesis, we have shown above that *Ac. thailandicus* can be continuously transmitted to the environment (Fig 6C, S10B Fig). Moreover, we detected bacteria in the surface of twenty out of twenty eggs laid by flies monoassociated with *Ac. thailandicus*, by testing bacterial growth in medium. This demonstrates that *Ac. thailandicus* is efficiently transmitted from mothers to their progeny.

We also observed that *Ac. thailandicus* affected the fertility of *D. melanogaster* in this assay. Similarly to the results above, there is no difference in total number of progeny if either or both parents are from the monoassociated stock (pupae or adult number, lm, *p* > 0.180 for all these comparisons). However, if both parents are axenic the number of pupae or adults total progeny is lower (*p* < 0.001 for all comparisons). This lower number of pupae or adults is not rescued by adding *Ac. thailandicus* to the axenic eggs (*p* = 0.998), indicating that these bacteria are not affecting egg to pupae or adult survival. Since exposing axenic adults to *Ac. thailandicus* does not alter their fertility (Fig 6D, 6E), this fertility effect may be dependent on either parent development in the presence of *Ac. thailandicus* or in the presence of *Ac. thailandicus* in the fly food for the two days of the egg laying.

The results above suggest that a stable association with gut bacteria is beneficial to adult *D. melanogaster*, because it allows continuous transmission to the next generation, promoting its faster development and higher fertility. Unstable interactions lead to loss of the bacterial population and non-transmission to the next generation. However, these experiments were performed by providing axenic food to flies, and in a natural scenario flies are bound to encounter many other bacteria present in the food substrates. If all bacteria were equally beneficial for fly development this stable association could be irrelevant. Therefore, we tested if different bacteria naturally encountered by *D. melanogaster* confer different fitness benefits to the flies. We sterilized eggs of *w^1118^ iso* and associated them with different bacteria found in the gut of flies from a natural population (sampled from the isolates of Fig 2, Fig 7A). We determined total number of adults that developed from these eggs, their developmental time, and their fertility. The number of adults that emerged (G0) was not different between associations with different bacteria or in germ-free conditions (S12A, S12B Fig, lmm, *p* > 0.282 for all pairwise comparisons). However, we did observe differences in the developmental time and fertility of these adults associated with different bacterial isolates, and found a negative correlation between these parameters (Pearson correlation −0.91, *p* < 0.001) (Fig 7B, S12C-S12F Fig, S13 Fig). Flies associated with *Ac. thailandicus* developed faster than axenic flies and flies associated with 11 out of the other 15 bacteria (lmm, *p* < 0.038 for all these pairwise comparisons). These flies are also more fertile than axenic flies and flies associated with 11 out of the other 15 bacteria (lmm, *p* < 0.018). Flies associated with *Ac*. OTU2753, *Lactobacillus brevis*, and *Lactobacillus paraplantarum* developed as fast and are as fertile as *Ac. thailandicus* (*p* > 0.200 for these pairwise comparisons). While flies associated with *Ac. cibinongensis* developed slower than with *Ac. thailandicus (p* = 0.023), the developmental time of flies with *L. pseudomesenteroides* is not significantly different (*p* = 0.224). However, both have lower fertility than flies with *Ac. thailandicus (p* < 0.001). On average, flies associated with *L. pseudomesenteroides* or *Ac. cibinongensis* develop faster and have a higher fertility than axenic flies but these differences are not statistically significant (*p* > 0.082, for all these comparisons). On the other hand, flies associated with *Bacillus flexus* OTU1589 were not different from axenic flies in terms of developmental time or fertility (*p* = 0.878). Overall, these data demonstrate that different bacteria have a variable effect on the development and fertility of *D. melanogaster*, with some not conferring any advantage to the flies development or fertility. *Ac. thailandicus* seems particularly beneficial to *D. melanogaster* and, therefore, the stable association may be advantageous to the host.

**Fig 7.**
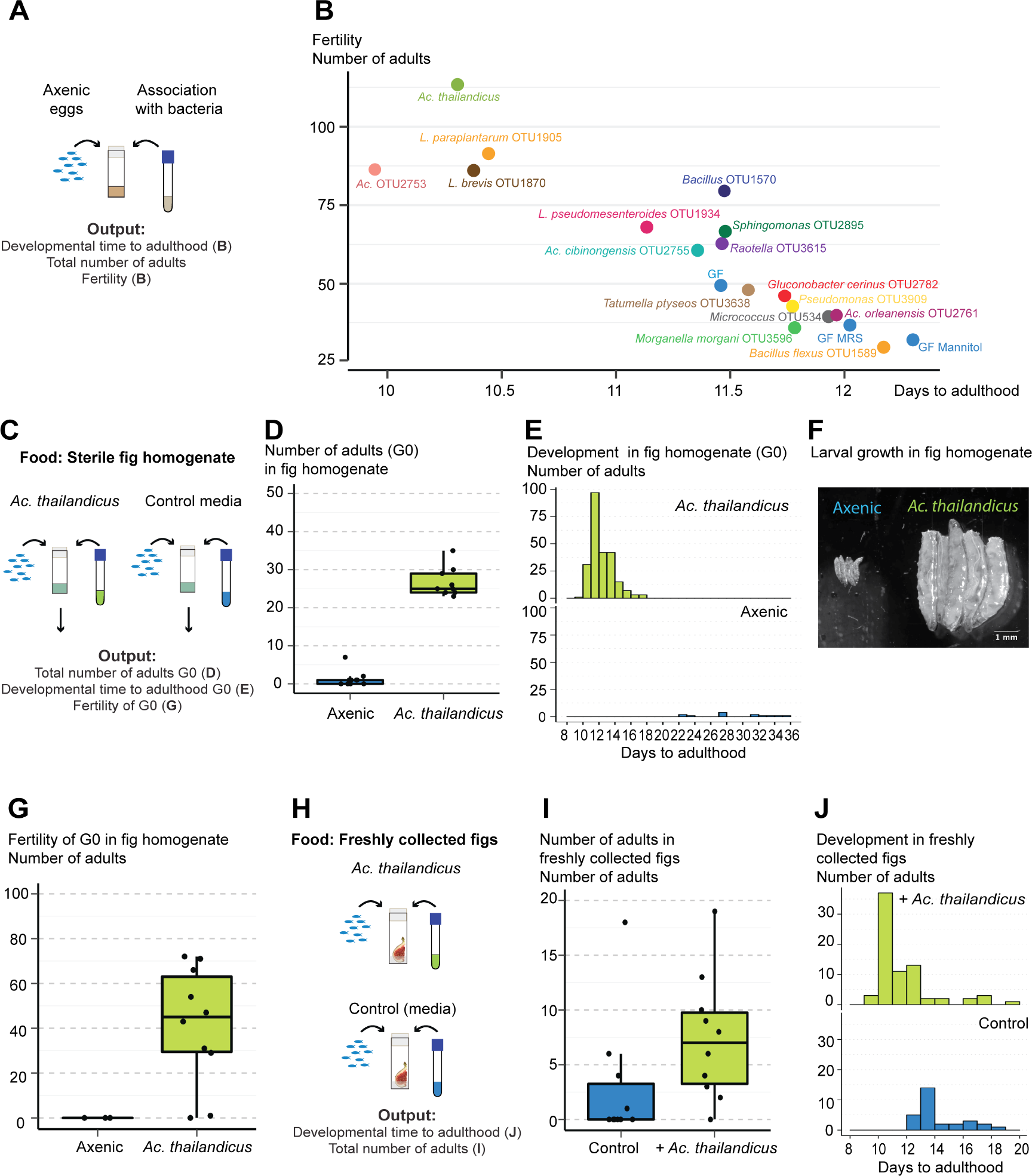
*Ac. thailandicus* is beneficial in the context of other wild bacteria and natural food substrates. (A) *w^1118^ iso* eggs were associated with different bacteria isolated from the gut of wild-caught *D. melanogaster*. As controls, axenic eggs that had no treatment (GF) or in which sterile media were added (GF MRS and GF Mannitol) were used. (B) For each bacterium, estimates of developmental time to adulthood of these eggs are plotted against estimates of their fertility. These estimates derive from the statistical analysis of data presented in S12C-F and S13 Fig. There is a negative correlation between developmental time and fertility (Pearson correlation −0.91, *p* < 0.001). (C) Fifty axenic *w^1118^ iso* eggs were placed in vials containing sterilized fig homogenate. *Ac. thailandicus* or sterile culture media were added on the top of the eggs. Ten vials were used per condition. Total number of adults that emerged (D) and developmental time to adulthood (E) was determined. More eggs inoculated with *Ac. thailandicus* developed to adulthood and faster than axenic eggs (lmm, *p* < 0.001 for both comparisons). (F) Larvae five days post inoculation with either condition in fig homogenate. (G) Fertility of flies developed in fig homogenate with and without the addition of *Ac. thailandicus*. Two males and one female were collected from G0 and placed per vial containing fig homogenate for 10 days, with vials flipped every other day. The *Ac. thailandicus* condition has ten replicates but only three from axenic eggs were possible to perform. Adults from eggs inoculated with *Ac. thailandicus* were more fertile than axenic adults (lmm, *p* = 0.003). (H) Fifty axenic *w^1118^ iso* eggs were placed in vials containing freshly collected non-sterile figs. *Ac. thailandicus* culture or sterile media (Control) was added on the top of the eggs. The total number of adults that emerged (I) and their developmental time to adulthood (J) was analyzed. Ten vials were analyzed per condition. There were more adults emerging from vials inoculated with *Ac. thailandicus* (lmm, *p* = 0.010). Developmental time to adulthood was faster in eggs inoculated with *Ac. thailandicus* in this experimental replicate but not significantly different in the other replicate represented on S14E Fig (lmm, *p* < 0.001 and *p* = 0.557, respectively). Statistical analyses from (D-J) were done together with replicate experiments shown in S14.

We also analyzed the impact of *Ac. thailandicus* on *D. melanogaster* fitness when they develop in fruit, a more natural food substrate, instead of standard fly food. We compared development from eggs to adults on a sterile fig homogenate with or without adding *Ac. thailandicus* (Fig 7C). The association with *Ac. thailandicus* strongly influenced the number of emerging adults, with very few flies reaching adulthood in axenic conditions (Fig 7D, S14A Fig, lmm, *p* < 0.001). Moreover, while *Ac. thailandicus* associated flies develop, on average, in 11.5 days, the few axenic flies that reach adulthood are slower and take 28 days (Fig 7E, S14B Fig, lmm, *p* < 0.001). This reflects a delay in growth since five days old larvae in axenic conditions were much smaller than larvae with *Ac. thailandicus* (Fig 7F). We subsequently tested the fertility of the adult flies that developed in these two conditions. Adults that developed on figs in the presence of *Ac. thailandicus* were also more fertile (Fig 7G, S14C Fig, lmm, *p* = 0.003). In fact, the few flies that developed in axenic conditions were all sterile. These results show that *Ac. thailandicus* benefit for the development and fertility of flies is even more pronounced in a natural food substrate.

However, in nature, fruits are not sterile but exposed to many environmental organisms. Therefore, the advantage we observed of *Ac. thailandicus* could be absent in normal non-sterilized fruit. Thus, we further tested the potential benefit of *Ac. thailandicus* by comparing the development of *D. melanogaster*, from axenic eggs, in non-cleaned, freshly collected figs, in the presence or absence of these bacteria (Fig 7H). Flies grown in the presence of *Ac. thailandicus* had approximately the double of the survival rate to adulthood than control flies with no bacteria added (Fig 7I, S14D Fig, lmm, *p* = 0.010). This is similar to the effect seen in sterile figs. The effect of *Ac. thailandicus* on the time to reach adulthood varies with replicate (Fig 7J, S14E Fig, lmm, *p* < 0.001). In one replicate the bacteria presence does not affect time of development (S14E Fig, *p* = 0.557), while in the other replicate *Ac. thailandicus* decreases time of development by 3.5 days (Fig 7J, *p* < 0.001). This difference between replicates may reflect the variable bacteria consortiums in the figs collected at different times. These results support that the stable association between *D. melanogaster* and *Ac. thailandicus* is beneficial for the flies in their natural environment.

## Discussion

Here, we identify bacterial isolates from a natural population of *Drosophila melanogaster* that can proliferate and stably colonize the gut of their host. These results demonstrate that *D. melanogaster* has *bona fide* gut bacterial symbionts in the wild. We further show that the association with one of these gut bacterial symbionts, *Acetobacter thailandicus*, can be mutually beneficial. On one hand, stable colonization of *D. melanogaster* gut permits continuous bacterial shedding to the environment, and, therefore, potentially increasing bacterial dispersion in the wild. On the other hand, transmission of *Ac. thailandicus* to the food substrate, concomitant with egg laying, benefits *D. melanogaster* larval development. These bacteria shorten developmental time and increase fertility of *D. melanogaster*. This stable interaction may be particularly important for *D. melanogaster* since different bacteria affect differentially its development and *Ac. thailandicus* is more beneficial than most bacteria sampled from the gut of wild flies. Moreover, *Ac. thailandicus* is still beneficial when larvae develop in non-sterile fruit collected from nature.

### Diversity and stability of gut bacteria in wild and laboratory *D. melanogaster*

In this study, one of our main concerns was to quantify, in different scenarios, absolute levels of live bacteria in the gut of *D. melanogaster*. Therefore, the several protocols we developed were mainly based in culture dependent techniques. This approach also allowed us to isolate bacteria for further functional characterization. Moreover, gut bacteria of *D. melanogaster* previously identified through 16S rRNA gene sequencing [20,26,28,41-46] belong to genera that can also be identified by culture dependent techniques. However, it is possible that our approach missed gut bacteria that do not grow in the media or conditions that we used. Additionally, our approach mainly identifies the bacterial strains that are more frequent in the gut as there is a limited number of colonies in the plates analyzed. Because of these limitations our analysis may be incomplete. Nonetheless, our approach managed to quantify overall gut bacterial numbers in different husbandry conditions, and, when tested, the results were confirmed by quantitative PCR. Moreover, we were able to identify, isolate, and analyze bacteria that can stably associate with *D. melanogaster* gut.

Our results show a striking difference in gut bacterial diversity between lab and wild caught flies. Lab flies carry mainly two bacterial species corresponding to *Acetobacter* OTU2753 and *Lactobacillus* OTU1865. This low diversity and dominance of *Acetobacter* and *Lactobacillus* species is in agreement with several previous studies on the gut associated bacteria in lab flies [20,22,24-27]. On the other hand, wild caught flies have a much higher diversity of bacteria. We were able to identify 35 different OTUs in the ten individual flies freshly collected from the wild, and the sampling did not seem close to saturation. This higher diversity is also in agreement with previous reports [25,28]. The characterization of individual flies allowed us to identify *Enterobacteriaceae, Acetobacteriaceae* (mainly *Acetobacter* and *Gluconobacter* species), *Leuconostocaceae*, and *Bacillaceae* as the most prevalent families, present in over 50% of the flies. These families of bacteria have been identified before in wild caught *D. melanogaster*, although *Bacillaceae* are found less frequently [25,28,41-43,46]. *Lactobacillus* was found in only one out of ten freshly collect wild flies analysed. Although the low prevalence of *Lactobacillus* could be a characteristic of this specific population, it is a general trend observed in other published surveys [25,28,41-43,46].

We tested persistence of bacteria in the gut of *D. melanogaster* by regularly changing individual flies to fresh axenic food and, therefore, reducing the potential intake of bacteria from contaminated food. This protocol is alike the one used in Blum et al. 2013 [20]. Similarly to that paper, we also found that the *Acetobacter* and *Lactobacillus* species associated with this laboratory stock cannot stably persist in the gut. Moreover, we show that these bacteria can grow in the fly food. Thus, these bacteria are only transiently passing through the gut. This result highlights how husbandry conditions can affect *D. melanogaster* gut bacterial levels and that these measured levels can be unrelated with gut colonization (also shown in [20,27]).

In contrast to lab flies, wild caught flies carry bacteria that, following this protocol, persist in the gut of *D. melanogaster*. This shows that in its natural state *D. melanogaster* lives with gut colonizing bacteria. *L. pseudomesenteroides, Ac. cibinongensis* and *Ac. thailandicus* were each present in more than 50% of wild flies at the end of the stability protocol. They are, therefore, interesting bacteria to further characterize in their interaction with *D. melanogaster*.

Several bacteria were present in 50% or more of the flies when they were caught, but were severely reduced in frequency after the stability protocol. These include *Bacillus* OTU1570, the *Enterobactereaceae* OTU3529, *Tatumella* OTU3635, and *Kluyvera ascorbata* OTU3643, and the *Acetobacteraceae Ac*. OTU2753, *Ac. ghanensis* OTU2757, *Ac. lovaniensis* OTU2759, *Ac. orientalis* OTU2760, and *Gluconobacter* OTU2781. These species may be transient gut bacteria that were acquired from the environment. However, it is also possible that they are stable gut bacteria that cannot be sustained in the particular lab environment we used. For instance, in the fly food we used there may be nutritional requirements missing for their maintenance or there could be toxic compounds to them (e.g. methylparaben). In the future, this protocol could be repeated using other food source, as for example the fruit matching the source of capture. However, it will be difficult to assert that a particular bacterial strain cannot persist in the gut even if it fails to show that property under more natural conditions. The natural environment of *D. melanogaster* is very complex and includes decomposing and fermenting fruit replete with different microorganisms. This will be hard to replicate and study in a controlled lab setup.

At the end of the stability protocol there was still a high diversity of bacteria in the gut of *D. melanogaster* even if most were present in less than 50% of the flies. These may represent rare but stable gut bacteria of *D. melanogaster*, as the case of *Lactobacillus* species. A particular fly (fly 39 in Fig 2) has an interesting pattern of microbiota composition after the stability protocol. It is the only wild caught fly that has no *Lactobacillales* or *Acetobacteriaceae*. Instead it carries six rare OTUs at relatively high levels. This gut microbiota composition may represent a disease-related dysbiosis and some of these bacteria could be pathogenic.

### Gut colonization *by Ac. thailandicus* and *Ac. cibinongensis*

To further characterize the interaction of the bacteria that persist in the gut of wild flies, we studied them in monoassociation with lab flies. In contrast to the lab isolate of *Ac*. OTU2753, both *Ac. cibinongensis* and *Ac. thailandicus* persist in the gut of lab flies until the end of the stability protocol. However, the levels of both bacteria decreased significantly in the first two days of this assay. These results indicate that the majority of the bacteria found in the gut of these flies at the beginning of the experiment were transient and lost with the same dynamics as unstable bacteria, but a certain part of these two bacterial populations are stably associated with the host. These results, in monoassociation, demonstrate that either of these bacteria have the autonomous property to persist in the host, independently of other microbiota members. Moreover, this property seems largely independent of host background since it is observed in the *w^1118^ iso* lab flies and in several individuals of the natural outbred population.

Both *Ac. thailandicus* and *Ac. cibinongensis* are able to proliferate in the gut of *D. melanogaster*. Interestingly, *Ac. thailandicus* seems to proliferate faster and reaches higher levels in 24h, which is coherent with higher bacterial levels in the stability protocol. The stability and proliferation assays show that these bacteria are *bona fide D. melanogaster* gut colonizers.

The niche of the stable population of *Ac. thailandicus* is the anterior gut of *D. melanogaster* since it is present in the crop and anterior midgut samples and absent from the mid midgut to the hindgut. The crop is a diverticulum of the oesophagus that can store liquid food [47]. In our analysis, the anterior midgut sample also included the proventriculus, which is part of the foregut. This raises the possibility that *Ac. thailandicus* stable population is restricted to the foregut. The epithelium in the foregut region has a cuticular lining, which could provide a surface for the bacteria to attach. Also, the crop lumen is not subject to the same linear flux as the rest of the gut lumen, which might facilitate bacterial persistence. A similar argument is made for the appendix and cecum, in humans and other mammals, as a reservoir of microbiota [48,49].

The midgut has a different structure and kinetics to the foregut. In the midgut the peritrophic matrix separates the gut lumen from the epithelial cells. This barrier is continuously secreted at the proventriculus and moves through the midgut with food [50,51], which may hamper stable bacterial colonization. Moreover, the foregut-midgut border may work as a physical or immunological barrier for microorganisms in insects [27,52-54], and the acidic region in the anterior midgut may also contribute to bacteria killing [38,55]. A reduction in bacterial loads after this transition is evident for *Ac. thailandicus* even before the stability protocol, and was previously observed in *D. melanogaster* gut [27,38].

The anterior gut may be a common location for bacterial colonization in *D. melanogaster. Pseudomonas aeruginosa*, a pathogenic bacteria, also colonizes the crop, where it forms a biofilm [56], and it was also suggested that the anterior gut could be a site for stable attachment of *Lactobacillus plantarum* [31]. Moreover, the crop was identified as the region where yeasts proliferate in flies, 130 years ago [57]. In the future it will be interesting to investigate where other *D. melanogaster* bacterial gut colonizers reside (e.g. *Ac. cibinongensis, L. brevis)*. In other insects gut bacteria are known to colonize diverse locations, including the proventriculus, the posterior midgut, and the hindgut [58-63].

### *D. melanogaster* and *Ac. thailandicus* mutualism

Given the stable association between *D. melanogaster* and *Ac. thailandicus*, we asked if there was any advantage for either partner in this interaction. Symbiosis between a host and a microbe does not necessarily signifies mutualism and the effect of host-association on the microbial partner has been less frequently studied [64,65]. Our results indicate that the stable association of *Ac. thailandicus* to the gut of the adult fly is advantageous to this bacterium since it can promote its dispersal.

The interaction with *Ac. thailandicus* is also advantageous to *D. melanogaster* in several scenarios. *Ac. thailandicus* shortens larvae developmental time to pupariation and adulthood when compared to axenic conditions. This effect does not necessarily increase fitness but it may, if there are no associated trade-offs, as shown with *L. plantarum* [37]. Interestingly, adult flies that developed in the presence of *Ac. thailandicus* are also more fertile, a clear measure of fitness, when compared with flies that developed axenically. These phenotypes demonstrate the benefit of *Ac. thailandicus* during *D. melanogaster* development. Other bacteria have been shown before to shorten development time of *D. melanogaster* [15,16,66-69] and increase adult fertility when associated in larval stages [70]. Moreover, adding the unstable lab isolate *Ac*. OTU2753 to axenic eggs also had a similar effect to adding *Ac. thailandicus*, in terms of developmental time and later adult fertility. So the direct developmental benefit conferred by these *Acetobacter* does not seem dependent on the capacity to colonize the gut of *D. melanogaster*. However, the most interesting aspect of this result is that, out of the 15 bacteria isolated from wild flies, *Ac. thailandicus* induced the shorter development time and higher fertility. Therefore, out of the set of bacteria interacting with *D. melanogaster* in the wild, this stable gut symbiont is particularly beneficial.

We do not know the mechanism through which *Ac .thailandicus*, or the other bacteria we tested, benefit *D. melanogaster*. The negative correlation that we observed between developmental time and fertility, suggests a similarity on the mechanisms behind these phenotypes. Microorganisms have for long been recognized as important for *Drosophila* development and as a source of food [14,71]. In fact the standard *Drosophila* food used in the lab is partly composed of dead *Saccharomyces cerevisiae* [72], which, in this diet, is required and sufficient for *Drosophila* development. Moreover, in lab diets the bacterial influence on host development is generally stronger the less yeast extract the food contains [15,16]. A recent study with *L. plantarum* also shows that heat-killed bacteria can rescue growth in germ-free conditions almost to the same extent as live bacteria [38]. In adults, constant supply of heat-killed yeast *Issatchenkia orientalis* can also extend the lifespan of *D. melanogaster* to the same extent as live yeast [19]. The nutritional value of these microorganisms may be based on supplying aminoacids or vitamins to the host [14,19,71,73]. Other evidence indicates that the effect of microorganisms on development of *D. melanogaster* could also be independent of its nutritional value. Bacteria can directly impact host physiology by activating the insulin pathway, via acetic acid production in the case of an *Acetobacter pomorum*, or gut proteases in the case of *L. plantarum* [16,39,74].

The benefit of *Ac. thailandicus* for *D. melanogaster* becomes even more evident when larvae develop in figs, a natural food substrate. On sterile figs homogenates very few larvae reach adulthood in axenic conditions, and those that do are severely delayed in growth and are infertile as adults. These results show the insufficiency of fruit, or figs in this particular case, to support normal *D. melanogaster* development. *Ac. thailandicus* rescues these phenotypes and is, therefore, sufficient for *D. melanogaster* development on fruit, indicating a nutritional basis for the interaction.

An alternative hypothesis is that bacteria are detoxifying some toxic components present on the food. Detoxifying symbiosis is known to occur in many insects [75]. However, the fact that *Ac. thailandicus* is beneficial both in lab food and figs indicate that to a large extent its benefit is independent of food toxins.

Although we saw that *D. melanogaster* benefits when it develops with *Ac. thailandicus*, we did not see a direct effect when flies are exposed to the bacteria only during adulthood. When we associated this bacterium to axenic adults, and they maintained a stable bacterial population for several days, their fertility did not change. However, direct effects of bacteria on adults have been previously reported on oocyte development or fertility [70,76]. Many factors may explain the different results, including the identity of the bacteria tested. Another explanation could be that the relatively small bacterial stable population in the gut, as in our assay, does not have an impact on host fertility, but higher levels of *Ac. thailandicus* would. The positive effect of *Ac. thailandicus* on the progeny of adults seem to be only due to being transmitted to the next generation and not to any effect on the adult itself. This is demonstrated by the fact that adding *Ac. thailandicus* to axenic eggs has the same effect, in terms of development, as having parents associated with the bacterium. Nonetheless, it will be interesting in the future to determine if the stable *Ac. thailandicus* population has any other effect on the adult physiology.

### Gut colonization by *Leuconostoc and Lactobacillus*

Analysis of *L. pseudomesenteroides* stability and proliferation in *D. melanogaster* gut produced ambiguous results. This bacterium seemed very stably associated with the gut of wild and monoassociated lab flies when the stability protocol was performed in vials. When we implemented the protocol using cages, however, it disappeared from 50% of the flies. These results illustrate how sensitive to experimental conditions is this assay, and that stringency is crucial. The proliferation assay did not clearly show an increase or decrease in *L. pseudomesenteroides* at 24h, when compared to the beginning of experiment. These results could be the consequence of this bacterium being able to very rapidly proliferate in the gut of the fly but unable to attach to the host and, therefore, require a constant cycle of re-inoculation. Maybe this cycle could be kept in vials but broke down in cages. Further experiments will be required to test this hypothesis and elucidate the interaction of *L. pseudomesenteroides* with *D. melanogaster*.

*Lactobacillus* species were still present in wild flies at the end of stability protocol, although less frequently than *Ac. thailandicus, Ac. cibinongensis*, and *L. pseudomesenteroides*. Interestingly, the data indicate a negative interaction between *Lactobacillus* and *Leuconostoc* presence. Both are lactic acid bacteria (order *Lactobacilalles)* and they may occupy the same niche and compete for resources. Of the many bacterial isolates from the gut of wild flies, *L. brevis*, and *L. paraplantarum* are the most beneficial in terms of development time and fertility of *D. melanogaster*, together with *Ac. thailandicus*. This contrasts with previous reports indicating a small or null effect of lab *Lactobacillus* isolates on fecundity [37,70]. *L. brevis* is present in four out of ten wild flies after the stability protocol and proliferates in the gut of *D. melanogaster*. So, *L. brevis* may also be a beneficial *bona fide* gut symbiont of *D. melanogaster*, although not as frequent as *Ac. thailandicus* in this population.

### Ecological advantage of a stable gut association with beneficial bacteria

Our results indicate that the interaction between *D. melanogaster* and the gut symbiont *Ac. thailandicus* is especially beneficial for both partners in the wild (Fig 8). The small stable bacterial population in the gut serves as a reservoir for the inoculation of the environment that the adult fly explores and exploits. This is beneficial to the bacteria since it leads to their continuous dissemination. On the other hand, transmission of *Ac. thailandicus* to the food substrate of the next generation, concomitant with egg laying, benefits *D. melanogaster* development. This association is therefore a form of farming, a strategy adopted by several insects, including ants, termites and ambrosia beetles with fungi [77]. The stability of the *D. melanogaster–Ac. thailandicus* interaction provides the host some independence from the local bacterial populations and enables it to explore and modulate bacterial populations in new locations.

**Fig 8.**
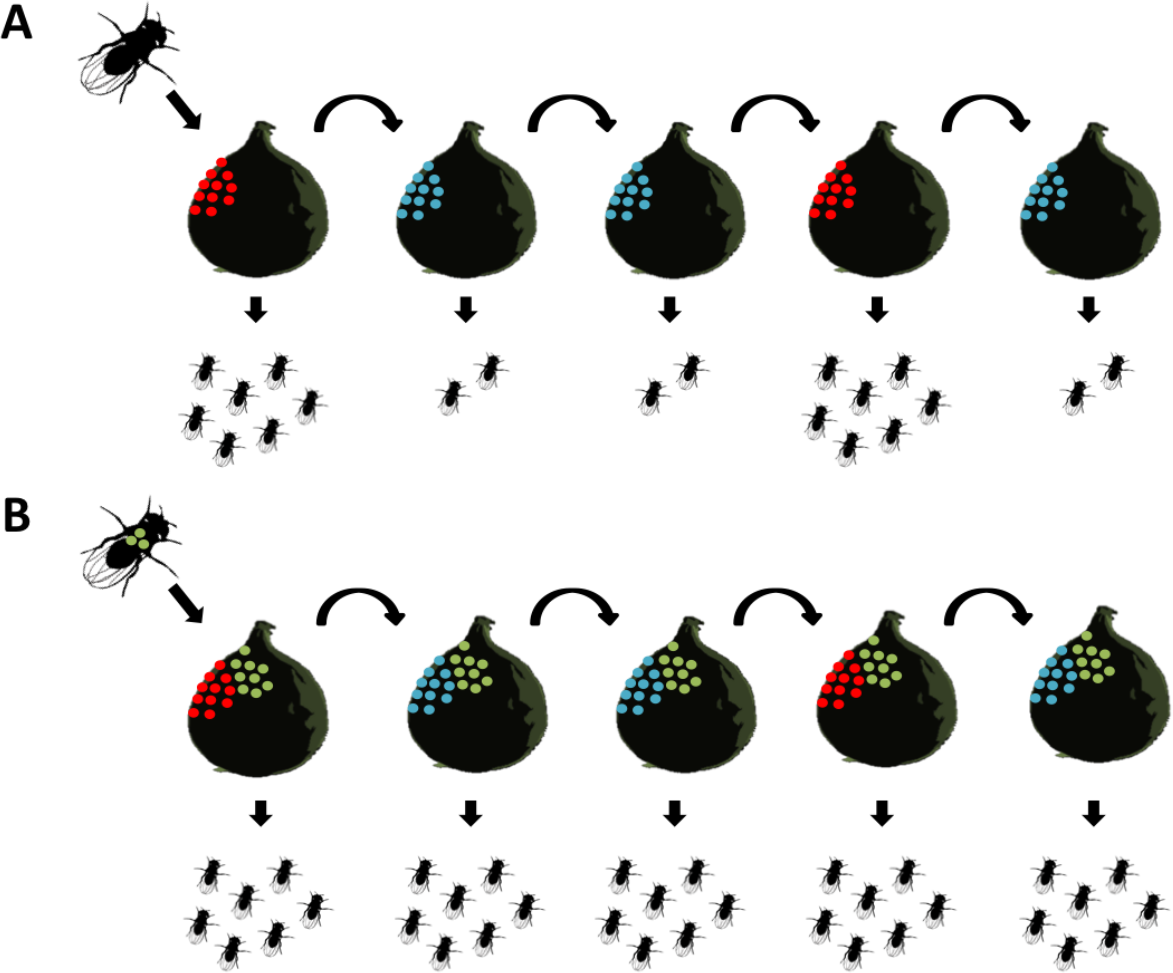
Model for an ecological advantage of a stable association between *D. melanogaster* and beneficial gut bacteria. (A) In the absence of stable gut bacteria, the fitness of *D. melanogaster* is dependent on the presence of more (red) or less (blue) beneficial bacteria in the food substrate. (B) Carrying a stable population of beneficial bacteria (green) in the gut allows constant bacterial inoculation of food substrate and consequent association with the next host generation. This leads to a higher fitness of this next generation.

Besides the interaction with these stable bacteria in the wild, *D. melanogaster* also interacts with a plethora of environmental bacteria that are transiently associated with the gut. Many of these non-colonizing bacteria probably positively impact on *D. melanogaster* biology, and vice-versa. *D. melanogaster* are attracted to feed on, or oviposit in substrates with specific potential benefiting bacteria [76,78-81]. Attraction to fermenting fruits enriched with beneficial bacteria may be a strategy adopted by *D. melanogaster* to increase interactions with these bacteria. Furthermore, *D. melanogaster* most likely disperses bacteria that transit through its gut. By attracting flies certain bacteria could, therefore, increase their probability of being dispersed. However, if bacteria are not stably associated with the flies, this would be a transient phenomenon, as evident in the rapid loss of *Ac*. OTU2753 in our experimental system involving a short exposure to bacteria and continuous change of food in cages. *D. melanogaster* may also benefit bacteria by promoting their growth in the food substrate [38], which could be advantageous for the host if biased towards beneficial bacteria. Despite all these potential mechanisms promoting beneficial interactions, relying on the immediate environmental and local bacterial community may be suboptimal for *D. melanogaster* (Fig 8).

*Ac. thailandicus* belongs to the acetic acid bacteria, a group of bacteria that oxidise the ethanol present on fermenting fruits to acetic acid. These bacteria are found associated with many *Drosophila* species and a wide range of other insect species, which normally rely on high-sugar diets [82,83]. Several *Drosophila* species are attracted by acetic acid bacteria and this is probably related with the production of acetic acid [79,80,84]. In addition, the aerobic environment and acidic pH of digestive tracts of most insects are suitable for acetic acid bacteria growth [82,85-87], and these bacteria produce extracellular matrixes, which can be involved in host adherence [83,88]. *Ac. thailandicus* interaction with *D. melanogaster* is another contribution to the understanding of the association of this group of bacteria with insects in an ecological context.

In the future it will be interesting to address some questions relevant for this model. For instance, we do not know how stable is *Ac. thailandicus* in the gut of larvae or if this stability is important. It may be sufficient for the bacteria to grow on the food substrate since larvae are less mobile and they will be in constant contact with the local external population of bacteria. Another important aspect is to understand how adult flies acquire *Ac. thailandicus*. This could be through constant association throughout the developmental stages, including from larvae to pupae to adult, or *de novo* acquisition after adult eclosion [66].

This farming interaction model may extend to other bacteria, including *L. brevis*. Moreover, our study focused on the gut colonizing bacterial species in one *D. melanogaster* population. It will be important to analyze other natural populations and determine to what extent there is conservation of stably colonizing species or if different *D. melanogaster* populations harbor different gut bacterial symbionts. This analysis could elucidate if there is a core gut microbiota of *D. melanogaster* based on stable symbionts. Diet has been shown before to influence the composition of the total bacteria associated with *D. melanogaster* [25,28,89]. Thus, it would be interesting to investigate how diet, or geography, determines the stable gut bacterial community. Moreover, additional studies need to be performed to identify other types of microbes that can stably associate with *D. melanogaster*. Particularly, it would be important to identify natural yeasts isolates that would colonize *Drosophila* intestine, given that flies are constantly exposed to different yeasts in the natural habitat.

Interactions between microbes may affect their colonization and their influence on host phenotypes. These may happen with other colonizing bacteria or with environmental bacteria on the food substrate or while in transit through the gut. Our analysis of wild-caught flies incorporates, to a certain degree, this complexity. For instance, *Ac. thailandicus* that stably colonizes in monoassociation is also present in the gut of the majority of wild flies of the population we analyzed, showing that its association is robust in the face of rich bacterial communities. Moreover, the beneficial effect of this bacterium observed in monoassociation is also present in the context of complex and natural microbial communities of figs. On the other hand, the analysis of wild-caught flies also indicates a negative interaction between *Lactobacillus* and *Leuconostoc* species.

### Specificity of gut symbionts

*Ac. thailandicus* can colonize the gut of *D. melanogaster* but not of *D. simulans*. On the other hand, *Ac. thailandicus and Ac. cibinongensis* seem to be the only stable *Acetobacter* species in the population we analyzed and *Ac*. OTU2753 from the lab cannot colonize the gut of *D. melanogaster*. This indicates that these stable interactions are specific from both host and symbiont perspectives. Subtle differences in the bacteria associated with *D. melanogaster* and *D. simulans* in the wild have been found before [28] but differences may be clearer when looking into the stable gut symbionts of different *Drosophila* species.

The presence of these species-specific mutualistic interactions of gut bacteria with *D. melanogaster* raises the possibility that these are long-term interactions and the result of adaptation. Therefore, they may be a good system to study host-symbiont evolution and even address questions of co-evolution and co-speciation [30,90-92].

We do not know the cause of the specificity of these colorizations. The interaction between the host immune system and different bacteria could be one of the mechanisms involved in this selection. Pathogenic bacteria can down-regulate or escape from the host immune system to establish infection [93]. In *D. melanogaster* alterations in immunity have an impact on gut bacterial compositions or load [22,27,94]. In mosquitoes the expression of host C-type lectins protects gut bacteria from antimicrobial peptides (AMPs) action in the gut and therefore modulate the gut bacterial community [95]. Many innate immune genes in *Drosophila* species are under fast positive selection [96-98] and differences in these genes could mediate association of different *Drosophila* species with different stable gut bacteria.

### Stable gut bacteria in *D. melanogaster* as an experimental system

Although the perspective of a transient microbiota has been dominant in most analyses of gut bacteria in *Drosophila* [20,27,38,43,99], there is some evidence of stable gut bacteria in these flies. Recently it was shown that a wild isolate of *Lactobacillus plantarum* has a higher frequency of gut colonization than a lab isolate [31]. These results are in agreement with a tendency for wild isolates of bacteria being better at colonizing *D. melanogaster*. However, in this study once bacterial colonization was established, titres were constant over time in wild and lab isolates [31]. It will be interesting to also test these isolates with the proliferation and stability protocols that we describe here. On a different approach, analysis of wild-caught individuals from other mushroom and cactus-feeding *Drosophila* species have identified bacterial strains highly enriched in the gut but very poorly represented in matched substrate samples [29,30]. This indicates that these enriched bacteria are gut symbionts and it will be also interesting to study them in more detail.

The presence of stable associations in the wild raises the question of why these seem to have been lost in laboratory stocks. Part of the answer may be related with the fact that association with non-colonizing bacteria can be as beneficial as with colonizing bacteria in the lab (e.g. *Ac*. OTU2753 vs *Ac. thailandicus)*. Fly husbandry conditions in the lab normally ensure transmission of bacteria from generation to generation even if they do not stably colonize the gut. Therefore, under laboratory conditions, there may be a loss of selective pressure for stability. This can lead to loss of the capacity to stably colonize the gut by the bacteria either by drift or by selection if there is a cost associated with this capacity. Alternatively, colonizing bacteria may be replaced by non-colonizing strains in the lab. The lab diet is relatively uniform and different from the natural diet, therefore, bacteria better adapted to these conditions may outcompete wild isolates [100]. Moreover, use of antifungal antimicrobials, and sometimes antibiotics, may constantly or occasionally severely disrupt bacterial communities associated with the flies that are then replaced with local bacterial strains that do not have the capacity to colonize *Drosophila*. One or combinations of these factors may over the long periods of time that flies are kept in the lab lead to the loss of the original microbiota. From our experience, wild bacterial isolates seem to be easily outcompeted in lab conditions and replaced by other bacteria, since we needed to carefully handle the fly stocks to keep the monoassociations with wild isolates.

Exploring the interactions between hosts and its natural colonizing symbionts can uncover new phenotypes missed in laboratory experiments. Previous studies with other organisms have shown that indeed this can be the case. For instance, in the nematode *Caenorhabditis elegans*, bacteria isolated from natural habitats conferred higher fitness when compared with the standard *E. coli* used in the laboratory [101,102]. Also, wild collected mice harbor a different microbiota to laboratory mice, which decreases inflammation and is protective upon infection and tumorigenesis [103]. The capacity to colonize and proliferate in the gut of *D. melanogaster* described in this study, demonstrates different properties from lab and wild bacterial isolates. Moreover, other phenotypes associated with this wild isolates may yet be identified.

The stable interaction we found between *D. melanogaster* and gut bacteria will be useful to address important questions in the gut microbiota field using this model system. This includes identifying and characterizing from the host and bacteria perspective genes required for colonization and for the control of this interaction. Moreover, it will allow understanding determinants of specificity, which are largely unknown, although adhesion and biofilm formation are important in this process [104,105]. These questions are also relevant to specifically understand better and manipulate insect gut symbionts. The release of insects with specific gut bacteria in interventions may be useful against pests (e.g. by increasing the fitness of sterile males [106]) and against vectors of disease (e.g. by increasing resistance to pathogens [107,108]. Knowing what regulates gut stability may be important for the success of these approaches.

Our work defines a new paradigm for the association between *D. melanogaster* and gut bacteria in which stable associations exist and contribute to the fitness of both partners in an ecological context. Therefore this new conceptual and experimental framework to study gut stable symbionts will contribute to the growing field of *Drosophila-microbe* interactions.

## Materials and Methods

### Wild fly collection, stocks source, and maintenance

Wild flies were collected with traps, with fallen figs as bait, placed for 24h under a fig tree in Oeiras, Portugal (GPS coordinates 38°41′32.1″N, 9°18′59.4″W). *D. melanogaster* and *D. simulans* males were identified according to [40]. All the material to collect and sort wild flies was sterilized prior to use.

DrosDel *w^1118^* isogenic stock (*w^1118^ iso*) [33] was used as a laboratory stock, unless otherwise indicated. The female lines *D. melanogaster* O13 and *D. simulans* O13 were established from single wild females collected in 2013, and latter identified to the species level. Other stocks used were *D. melanogaster* Canton-S (Bloomington Drosophila Stock Center at Indiana University, stock #1), and *D. simulans* A07 and J04 (Drosophila Species Stock Center from California University, stocks #14021-0251.260, and #14021-0251.187, respectively). Unless otherwise indicated flies were 3-6 days in the beginning of experiments. The age of wild-caught flies is uncontrolled.

Stocks were kept and experiments were performed at 25°C in standard *Drosophila* food composed of 1.05L water, 80g molasses, 22g beet syrup, 8g agar, 10g soy flour, 80g cornmeal, 18g yeast, and 30ml of a solution containing 0.2g of carbendazim (Sigma) and 100g of methylparaben (Sigma) in 1L of absolute ethanol. Food was autoclaved before dispensing it into vials.

### Bacterial culture

Analysis of bacteria present in the gut was performed by culture dependent methods in order to isolate bacteria for further manipulations. From each fly the gut (including crop, midgut, and hindgut) together with the Malpighian tubules was dissected in Tris-HCl 50mM, pH 7.5, and homogenized with a plastic pestle in an 1.5mL microcentrifuge tube with 250µL Luria Broth (LB). Each sample was serially diluted (1:10 factor) and 30µL from each dilution were plated in five different culture media: LB (GRiSP), MRS (Merck), Liver Infusion Broth (Becton Dickinson), Brain heart infusion (BHI) (Sigma-Aldrich) and Mannitol (3g of Bacto Peptone (Becton Dickinson), 5g of Yeast Extract (Sigma-Aldrich), 25g of D-Mannitol (Sigma-Aldrich), 1L of Milli-Q water). Plates were incubated at 25°C for six days and dilutions containing 30-300CFUs were used to count and isolate bacteria.

To analyze flies or food associated with only specific bacterial isolates samples were plated on specific media to grow the correspondent bacteria (Mannitol for *Acetobacter* and MRS for *Leuconostoc or Lactobacillus)*. Plates were incubated at 25°C for 4 days.

### Quantification, isolation and identification of gut-associated bacteria

For quantification of total bacteria in each gut sample we selected the data from the medium that presented the highest number of colonies.

For a detailed analysis bacterial colonies were assigned, in each culture medium plate, per sample, to distinct morphological types and determined their number. Two colonies of each morphological type, per culture medium plate, per sample, were re-streaked and, after growth, colonies were picked, dissolved in 500µL LB containing 15% glycerol (v/v) and frozen at −80°C.

To identify each bacterial isolate a PCR was performed to amplify the 16S rRNA gene. For most samples a bacterial colony, or part of it, was directly placed in the PCR reaction tube (colony PCR). In the few cases where amplification was unsuccessful by colony PCR, DNA extraction was performed with ZR Fungal/Bacterial DNA MiniPrep Kit (Zymo Research according to the manufacturer’s instructions). Primers used were: 27f (5’-GAGAGTTTGATCCTGGCTCAG-3’) and 1495r (5’-CTACGGCTACCTTGTTACGA-3’) with the following PCR conditions: 94°C for 4min; 30 cycles of 95°C for 30sec, 58°C for 1min, and 72°C for 2min; 72°C for 10min. PCR products were sequenced at Source Biosciences Sequencing Center. Sequences were trimmed to 800bp of each sequence including V2 to V4 hypervariable regions. These sequences were aligned against a core set aligned fasta file from Greengenes [34] using PyNAST [109], and classified into operational taxonomic units (OTUs) according to Greengenes taxonomy [34]. Sequences that matched *Ralstonia* OTU3005*, Novosphingobium stygium* OTU2886, and *Novosphingobium* OTU2881 were removed from the analysis since they were occasionally present on negative controls for PCR.

In most cases each morphological type corresponded to one OTU. However, three groups of bacteria had different OTUs commonly assigned to the same morphological type. Thus, these bacteria could not be distinguished, within their group, based on colony morphology. These groups are composed of bacteria belonging to the *Lactobacillus* genus, the *Acectobacteraceae* family (*Acetobacter* and *Gluconobacter* genera), or the *Enterobacteriaceae* family. The frequencies of the sequenced colonies from each group are represented in Fig 3 and S3 Fig.

To determine CFUs per gut for each OTU, or group of bacteria, the data from the medium that presented the highest number of colonies was selected.

Bacterial isolates used for phenotypic analysis (*Ac*. OTU2753, *Ac. thailandicus*, *Ac. cibinongensis, L. pseudomesenteroides* and all isolates used in Fig 7B and S12) were sequenced with both 27f and 1495r primers, and analyzed at least from V2 to V8 hyper variable regions of the 16S rRNA sequence. Sequences were automatically edited with PhredPhrap and consensus sequences were generated using BioEdit Sequence Alignment Editor Software. Sequences are in S1 text and deposited in GenBank with the following accession numbers: MG808351.1, MG808350.1, MG808352.1, MG808353.1, MG808354.1, MG808355.1, MG808356.1, MG808357.1, MG808358.1, MG808359.1, MG808360.1, MG808361.1, MG808362.1, MG808363.1, MG808364.1, MG808365.1.

### Real-time quantitative PCR for 16S rRNA gene

DNA was extracted from dissected single guts with QIAamp DNA Micro kit (Qiagen) as described in the protocol “isolation of Genomic DNA from Tissues”. To facilitate DNA extraction from Gram-positive bacteria the guts were homogenized in 180µL of enzymatic lysis buffer with Lysozyme (from DNeasy Blood & Tissue Kit, QIAGEN) and incubated for 1 hour at 37°C, before starting the protocol. DNA concentrations were determined with a NanoDrop ND-1000 Spectrophotometer. Quantitative-PCR reactions were carried out in CFX384 Real-Time PCR Detection System (BioRad). For each reaction in 384-well plate (BioRad), 6 µL of iQ SYBR Green supermix (BioRad), 0.5 µL of each primer solution at 3.6 mM and 5 µL of diluted DNA were used. Each plate contained three technical replicates of every sample for each set of primers. Primers used to amplify the 16 S rRNA gene were: 8FM (5’–AGAGTTTGATCMTGGCTCAG-3’) and Bact515R (5’-TTACCGCGGCKGCTGGCAC-3’) [110]. Primers used to amplify *Rpl32* were: Rpl32 forward (5′-CCGCTTCAAGGGACAGTATC-3′) and Rpl32 reverse (5′-CAATCTCCTTGCGCTTCTTG-3′). The thermal cycling protocol for the amplification was: initial 50°C for 2 min, denaturation for 10 min at 95°C followed by 40 cycles of 30 sec at 95°C, 1 min at 59°C and 30 sec at 72°C. Melting curves were analyzed to confirm specificity of amplified products. Ct values for manual threshold of were obtained using the program SDS 2.4 or with Bio-Rad CFX Manager with default threshold settings. 16S rRNA gene levels were calculated relative to Day 0 sample with the Pfaffl method [111] using *Drosophila Rpl32* as a reference gene.

### Generation of axenic and monoassociated flies

To develop axenic flies, embryos were sterilized with 2% sodium hypochlorite during 10 minutes, followed by 70% ethanol during 5 minutes and washed with sterile water. Embryos were placed in sterilized food vials and maintained in axenic conditions or monoassociated with 40µL of overnight bacterial culture of specific isolates. Monoassociated stocks were kept at 25°C and flipped every 20 days, using sterile gloves. We waited at least two generations in monoassociation before performing experiments.

### Analysis of bacterial stability in the gut

The gut stability protocol in vials was based on placing a single fly per vial, with a food surface of 3.8 cm^2^, and changing it twice a day to new vials. The stability protocol in cages was based on placing a single fly per cage with six petri dishes with a total fly food surface of 486 cm^2^, and changing them daily. Bacterial levels were analyzed in single guts.

To analyze the gut region where stable bacteria are present, individual guts were dissected into 5 different regions - crop, anterior midgut, mid midgut, posterior midgut and hindgut. The proventriculus was included in the anterior midgut sample. Each gut region from a single fly was homogenized, plated, and quantified as described above.

### Analysis of bacterial proliferation in the gut

The proliferation assay was based on providing an inoculum of bacteria to axenic male flies for 6h and measure gut bacterial levels, by plating, immediately at the end of this period (time 0h), and 24h later. Bacteria were grown in the Mannitol (*Acetobacter*) or MRS (*Leuconostoc* or *Lactobacillus*) liquid media in a shaker at 28°C overnight. Bacterial concentrations (cell/ml) were calculated based on OD600 using a spectrophotometer (SmartSpec 3000 from Biorad) using the formula OD1 = 5 x 10^8^ cell/ml. The inoculum was provided in vials by adding 180µL of bacterial solution in 2.5% sucrose to a round filter paper placed on top of the fly food. After the inoculation period, flies were placed singly in cages or in bottles (food surface: 486 cm^2^ and 28 cm^2^, respectively) for 24h. Bacterial levels were analysed in single guts by plating.

To confirm that the 24h data corresponded to bacteria growing in the gut and not bacteria growing on the fly food and in transit we added an axenic fly to the cage or bottle at time 0, in some experiments. Bacterial levels in the gut of these chaser flies were determined at time 24h, simultaneously with the co-habiting experimental fly.

### Analysis of bacterial proliferation on fly food

To analyze bacterial growth on food from bacteria associated with flies, conventionally reared 3-6 days old males were placed singly in vials for 24h, in order to contaminate the food with bacteria. After that period, flies were discarded and vials were incubated for 9 days at 25°C. Bacterial levels were determined after discarding the flies (Day 1) and after the 9 days of incubation (Day 10). Vials that never contained flies before were used as control vials and incubated also for 9 days (Day 10 control). 2.9g of top layer of food were homogenized in 10mL LB. This homogenate was plated in the five different media.

To analyze growth of *Acetobacter* species on the fly food, 3-6 days-old males monoassociated with the different *Acetobacter* were singly placed in vials with 4ml of fly food for 16 hours. After that period, males were discarded and bacterial levels were assessed at that time-point (Day 0) and after 1 or 5 days of incubating the vials at 25°C. All the food from the vial was homogenized in 4ml LB. Mannitol plates were incubated at 25°C for 4 days.

### Fitness parameters determination

To determine fitness parameters in monoassociated stocks (S9 Fig) one virgin female and three 0-3-days old males were placed per vial for 3 days and then discarded. Time to pupariation and to adulthood was daily assessed, as well as total number of pupae and adults.

To analyze fitness parameters of flies in a changing environment (Fig 6, S10 Fig), axenic 1-3 days old females and males were in contact for 6h with an inoculum of 10^5^ CFU/µL *Acetobacter OTU 2753, Ac. thailandicus* or with sterile Mannitol. After this period, one female and two males were placed per cage for 10 days. Each cage contained six bottles with food that were changed everyday (total food surface of 170cm^2^). Single gut bacterial loads were analyzed in females 0 hours and 10 days post-inoculation and in males 10 days after inoculation.

From each cage, all the six bottles were daily collected, number of eggs was counted and bottles were kept to daily assess adult emergence (Fertility G0 and Development of G1). Transmission of bacteria to the food was analyzed in bottles without eggs at days 1, 3, 5, 7, and 9. The food surface was washed with 1000µL of Mannitol and 100µl of this suspension was plated in Mannitol. As a control, food from axenic flies was also tested at days 1 and 9 and no bacteria were detected.

To analyze fertility of G1, bottles from day 9 and 10 from each condition were used to collect flies. One female and one male from the same condition were placed per vial and flipped to new ones every other day, during 10 days. Adult emergence was daily assessed to determine total number of adults (Fertility of G1).

To analyze if the benefit of *Ac. thailandicus* was dependent on the association with either parent we compared the four possible pairs of males and females from an axenic stock and a stock monoassociated with *Ac. thailandicus* (Fig 6H, S11). We placed one female and one male, both 1-2 days old, per vial and flies were passed to new vials every other day during 10 days. We also tested a condition in which 30µL of an overnight *Ac. thailandicus* culture was added to the progeny of axenic parents immediately after emptying it of parents. We daily assessed developmental time to pupariation and adulthood.

To analyze fitness parameters conferred by different natural bacterial isolates (Fig 7B, S12 Fig), 50 sterilized eggs were placed per vial and inoculated with 40µL of an overnight bacterial culture. All isolates were grown at 28°C in Mannitol, except *L. brevis, L. paraplantarum* and *L. pseudomesenteroides* that were grown in MRS. As controls we analyzed sterilized eggs associated with only Mannitol or MRS, or with no medium added. Developmental time to adulthood (Number of adults (G0), days to adulthood (G0)) was assessed. One male and female of the first adults emerging from each condition were placed per vial and flipped every other day during 8 or 10 days. Adult emergence was daily assessed to determine total number of adults (Fertility of G0).

To analyze the impact of *Ac. thailandicus* on fitness parameters in sterile figs homogenate (Fig7 C-G), 50 sterilized eggs were placed per vial and inoculated with 40µL of an overnight culture of *Ac. thailandicus* or sterile Mannitol. Adult emergence was daily assessed. For the analysis of this G0 fertility, one male and one female adults that emerged from these vials were placed per vial and flipped every other day during 8 or 10 days. Adult emergence was daily assessed to determine total number of adults (Fertility of G0). The fig food homogenate was produced with 300mL homogenized commercial frozen figs, 600mL water, and 4.8g agar. After autoclave, food was poured to each vial in sterile conditions, inside a laminar flow hood.

To analyze the fitness impact of *Ac. thailandicus* in fresh figs, we collected these at the same location where the wild flies were collected. Figs were cut in quarters and placed them in vials with sterilized agar (0.8% agar in water) at the bottom to fix the fig. Thirty sterilized embryos were placed on the top of these figs and inoculated with 40µL of an overnight culture of *Ac. thailandicus* or sterile Mannitol. Quarters originated from the same fig were distributed to the two conditions. Adult emergence was daily assessed. As a control, figs without the addition of eggs were kept and no flies emerged from those ones. Also, all flies that emerged from the experimental conditions had white eyes, confirming that they developed from the sterilized eggs and not from a possible contamination with wild flies present in the figs.

### Statistical analysis

The statistical analysis was performed in R [112] and graphs were generated using the package ggplot2 [113] and GraphPad. The script of all the analyses is provided in S2 Text, where details can be found.

Bacterial levels, number of eggs, pupae and adults, and time to pupariation and adulthood were analyzed using linear models *(lm)*, or linear mixed-effect models (*lmer* package *lme4* [114]) if there were random factors. Significance of interactions between factors was tested by comparing models fitting the data with and without the interactions using analysis of variance (*anova*). Models were simplified when interactions were not significant. Pairwise comparisons of the estimates from fitted models were analyzed using *lmerTest* [115], *lsmeans* [116], and *multcomp* [117] packages.

Timecourse analysis of bacterial stability in cages was performed fitting a non-linear least-squared model with the parameters of an exponential decay curve. Model simplification was achieved through analysis of variation *(anova)* and Akaike information criterion *(AIC)* of fitted models.

Bacterial levels in flies in the changing environment cage assay were analysed with the non-parametric Mann-Whitney test *(wilcox.test)* because some data points were high and not estimated precisely.

Bacteria transmission to bottles in the changing environment cage assay was analysed with a generalized linear mixed-effects *(lme4* package) with a binomial distribution.

Independence of *Lactobacillus* and *Leuconostoc*, or different *Acetobactereaceae*, presence in wild-caught flies was tested with the Pearson’s Chi-squared test *chisq. test*.

Correlation between developmental time and fertility of flies that developed associated with different bacteria was tested through the Pearson correlation *(cor. test)* of the means of these parameters.

## Acknowledgments

We thank Tiago Marques for advice on the statistical analysis and Gonçalo Matos for comments on the manuscript. We are also thankful to Liliana Vieira and the IGC fly facility for help in producing fly food.

## Supporting information

**S1 Fig - Wild-caught *D. melanogaster* have a stable gut microbiota.** Single 3-6 days old *w^1118^ iso* males were kept in the same vial during ten days (A) or exposed to a stability protocol by being passed to new vials twice a day (A, B). (A) Five individuals were analyzed at each day and total number of CFUs per gut determined by bacterial plating. Bacterial levels increase in the flies maintained in the same vials and decrease in the flies flipped to new vials twice a day (lmm, *p* < 0.001 for both). (B) Relative amount of 16S rRNA bacterial gene was measured by quantitative-PCR in five individual guts from each day, using the host gene *Rpl32* as a reference gene. Relative amount of 16S rRNA gene decreases between days (lmm, *p* < 0.001). (C, D) Bacterial levels from wild-caught flies on the day of collection (Day 0) and after 5, 10 or 20 days of the stability protocol. Bacterial levels on the flies significantly decrease with time (lmm, *p* = 0.004). Each dot represents an individual gut and the lines represent medians. Statistical analyses were performed together with replicate experiments shown in Fig 1.

**S2 Fig - Higher diversity of gut bacterial communities in wild-caught *D. melanogaster*.** Accumulation curve of the different bacterial OTUs present in wild-caught and laboratory flies before (Day 0) and after (Day 10) being exposed to the stability protocol.

**S3 Fig - Total levels and diversity of *Enterobacteriaceae* in wild-caught *D. melanogaster*.** (A) Levels of *Enterobacteriaceae* in the gut of wild caught flies before (Day 0) and after 10 days of the stability protocol (Day 10). Each dot represents one gut and lines represent medians. Levels of *Enterobacteriaceae* decrease between days (lm, *p* = 0.01). (B) Frequencies of sequenced colonies of *Enterobacteriaceae* for Day 0 and Day 10, represented as several in Fig 2. Numbers on the top of the bars correspond to the number of flies carrying that specific OTU, from a total of 10 flies.

**S4 Fig – *Leuconostoc pseudomesenteroides* stably associates with the gut of wild *D. melanogaster*.** Total *L. pseudomesenteroides* levels in the gut of wild caught flies in the day of collection (Day 0) and after 10 days of the stability protocol (Day 10). Levels of *L. pseudomesenteroides* are not significantly different between days (lm, *p* = 0.372). Each dot represents one gut and the line represents the median.

**S5 Fig - *Ac. thailandicus* and *Ac. cibinongensis* stably colonize the gut of *Drosophila melanogaster*.** (A-I) Single 3-6 days old *w^1118^ iso* males from monoassociated stocks with *Ac*. OTU2753 (A, E), *Ac. cibinongensis* OTU2755 (B, F), *Ac. thailandicus* (C, G), or *L. pseudomesenteroides* (D, H) were exposed to the stability protocol for ten days in vials (A-H) or five days in cages (E-H). Number of CFUs in individual guts was assessed by plating before and after five or ten days of the stability protocol. Ten flies were analyzed for each condition. *Acetobacter* OTU2753, *Ac. cibinongensis*, and *Ac. thailandicus* levels decrease between day 0 and 10 day in vials (lmm, *p* < 0.001 for all), but *Leuconostoc pseudomesenteroides* levels do not significantly change (*p* = 0.96). (I) Data from Fig 4 B-E was fitted to an exponential decay model that estimates the exponential decay rate, which corresponds to the rate of bacterial loss from the gut, and an asymptote, that corresponds to the levels at which the bacteria levels tend to stabilize after this loss. The rate of decay is the same for all the bacteria but there are differences between the asymptotes of all bacteria (contrasts of nonlinear least-square model estimates, *p* < 0.014), except between *Ac*. OTU2753 and *L. pseudomesenteroides (p* = 0.395). (J, K) Number of CFUs in each gut region from *w^1118^ iso* males monoassociated with *Ac. thailandicus* before (F) and after (G) five days of the stability protocol in cages. Statistical analyses were performed together with replicate experiments shown in Fig 4B-G.

**S6 Fig - *Ac. thailandicus, Ac. cibinongensis* and *L. brevis* proliferate in the gut of *Drosophila melanogaster*.** Three to six days old axenic *w^1118^ iso* males were inoculated for 6 hours with different concentrations of *Ac*. OTU2753 (A, E), *Ac. cibinongensis* OTU2755 (B, F), *Ac. thailandicus* (C, G), *L. pseudomesenteroides* (D), *L. paraplantarum* (H, I) and *L. brevis* (J). Bacterial levels were assessed by plating 0 and 24 hours post-inoculation. During this period males were singly placed in cages. In (G) axenic chaser males were placed in cages together with males inoculated with *Ac. thailandicus*. At 24 hours bacterial levels were assessed in both males. Bacterial levels between 0 and 24 hours decrease in flies inoculated with *Ac*. OTU2753 (lmm, *p* < 0.001), increase in flies inoculated with *Ac. cibinongensis, Ac. thailandicus*, and *L. brevis* (lmm, *p* = 0.024, *p* < 0.001, and *p* = 0.046, respectively) and do not significantly change in flies inoculated with *L. pseudomesenteroides* and *L. paraplantarum* (lmm, *p* = 0.158 and *p* = 0.65, respectively). Four to five males were used per condition, except in (B) where three males were used at one time-point and in (D) where two males were used on the inoculation 10^4^ CFU/µl. Each dot represents one gut and lines represent medians. Statistical analyses were performed together with replicate experiments shown in Fig 4J-Q.

**S7 Fig *- Acetobacter* species grow on the fly food media.** Single 3-6 days old *w^1118^ iso* males from a monoassociated stock with *Ac*. OTU2753 (A, D), *Ac. thailandicus* (B, E) or *Ac. cibinongensis* (C, F) were placed per vials for a period of 16 hours and then discarded. Bacterial levels on the food were determined by plating after discarding the flies (Day 0) and after one or five days of incubating these vials. Levels of *Acetobacter* on the food increase for all conditions between Day and Day 5 (lmm, *p* < 0.001). Five vials were used per condition. Each dot represents the bacterial levels on the food of one vial and lines represent medians.

**S8 Fig *- Ac. thailandicus* proliferates in the gut of *D. melanogaster* and not in *D. simulans*.** (A, B) Optimization of proliferation protocol in bottles. Axenic 3-6 days old *w^1118^ iso* were inoculated for 6 hours with different concentrations of *Ac.* OTU2753 (A) or *Ac. thailandicus* (B). Bacterial levels were assessed 0 and 24 hours post-inoculation. During this period males were singly placed in bottles (food surface of 28.27cm^2^) together with an axenic chaser male, from which bacterial levels were also assessed at 24h. Levels of *Ac.* OTU2753 decrease between days (lmm, *p* < 0.001). Levels of *Ac. thailandicus* increase when flies are inoculated with the lowest concentration (*p <* 0.001) and are maintained when flies are inoculated with the highest concentration (*p* = 0.426). (C-E) Axenic 3-6 days old *D. melanogaster* or *D. simulans* males were inoculated for 6 hours with 10^3^ CFU/µl (C, D) or 10^4^ CFU/µl (E) of *Ac. thailandicus*. Bacterial levels were assessed 0 and 24 hours post-inoculation. During this period males were singly placed in bottles. Three different genetic backgrounds for *D. melanogaster (w^1118^ iso, D. mel*. O13 and Canton-S) and for *D. simulans (D. sim*. J04, *D. sim*. O13 and *D. sim*. A07) were used. Bacterial levels in the gut increase in *D. melanogaster* and decrease in *D. simulans (p <* 0.001). Five individuals were analyzed for each condition and total number of CFUs per gut determined by plating. Each dot represents one gut and the line represents medians. Statistical analyses were performed together with replicate experiment shown in Fig 5.

**S9 Fig - Flies monoassociated with *Acetobacter* develop faster and are more fertile than axenic flies in a constant environment.** (A-D) Total number of pupae (A), total number of adults (B), developmental time to pupariation (C) and developmental time to adulthood (D) was analyzed in flies from a monoassociated stock with *Ac.* OTU2753 or *Ac. thailandicus*, or in axenic flies. One female and three males from each condition were placed per vials for three days and then discarded. Number of pupae or emerged adults was daily assessed. Ten vials were used per condition. Flies monoassociated with either *Acetobacter* species develop faster and have higher fertility than axenic flies (lm, *p* < 0.001). (A, B) Each dot represents the total progeny of one female.

**S10 Fig *- Ac. thailandicus* stable association with *D. melanogaster* is mutualistic.** Axenic 1-3 days old *w^1118^ iso* males and females (G0) were in contact with an inoculum of 10^5^ CFU/µl of *Ac.* OTU2753 or *Ac. thailandicus*, for 6 hours. Two males and one female were placed per cage, with 5 cages for each condition, during 10 days with daily changed food. (A) Bacterial levels in single guts of females 0 hours and 10 days post-inoculation and in males 10 days post-inoculation, analyzed by plating. Bacterial levels between the two time-points increased in females inoculated with *Ac. thailandicus* and decreased in females inoculated with *Ac.* OTU2753 (Mann-Whitney test, *p* < 0.001 and *p* = 0.048 respectively). (B) Presence of bacteria on the food collected from cages at days 1, 3, 5, 7 and 9 of the protocol, analyzed by plating. Filled rectangles represent presence of bacteria. *Ac. thailandicus* is transmitted to the food with higher frequency than *Ac.* OTU2753 (glm-binomial, *p* < 0.001). (C-F) Effect of bacterial association on the fitness of *D. melanogaster*. Total number of eggs laid by flies inoculated with different *Acetobacter (C)* and total number of adults that emerged from these eggs (D). Total number of eggs or adults is not different between conditions (lmm, *p >* 0.484). (E) Developmental time to adulthood of the progeny (G1) of flies inoculated with different *Acetobacter*. Developmental time to adulthood is faster in progeny from flies inoculated with *Ac. thailandicus* than in progeny from flies inoculated with *Ac.* OTU2753 (lmm, *p* < 0.001). (F) Fertility of G1. Two males and one female of G1 were placed per vial and flipped every other day for 10 days. Five couples were made per condition. Total number of emerged adults was analyzed. Fertility is higher in progeny from flies inoculated with *Ac. thailandicus* compared than in progeny from flies inoculated with *Ac*. OTU2753 (lmm, *p* < 0.001). Statistical analyses were performed together with replicate experiments shown in Fig 6B-G.

**S11 Fig - Both parents transmit the beneficial effect of *Ac. thailandicus* to their progeny.** Combinations of one male and one female 1-2 days old *w^1118^ iso*, either axenic or monoassociated with *Ac. thailandicus* (Bact.), were placed in vials and flipped every other day for 10 days. To one set of vials with axenic parents *Ac. thailandicus* was added on the eggs after passing the parents. Ten couples were made per condition. Developmental time to pupariation (A, E), to adulthood (C), total number of pupae (B, F) and total number of adults (F, G) was assessed. (A-D) correspond to one experimental replicate and (E-G) correspond to another experimental replicate, together with data from Fig 6H. Progeny from couples where either or both parents are monoassociated and progeny from axenic flies where *Ac. thailandicus* culture is added on the eggs develop faster than progeny from axenic flies (lmm, *p* < 0.001, for all these comparisons). Total number of progeny (pupae or adults) from couples where either or both parents are monoassociated with *Ac. thailandicus* is higher than in progeny from axenic flies (lmm, *p* < 0.001). (B, D, F, G) Each dot represents the total progeny of one female. Statistical analyses were performed together with replicate experiment shown in Fig 6H.

**S12 Fig - Different bacterial species have different impact on host developmental time and fertility.** Fifty *w^1118^ iso* eggs were associated with different bacteria isolated from the gut of wild- caught *D. melanogaster*. As controls, axenic eggs that had no treatment (GF) or in which sterile media were added (GF MRS and GF Mannitol) were used. Ten vials were used for each condition. Total number of emerged adults (A, B) and their developmental time to adulthood was daily assessed (C, D). Number of emerged adults is not significantly different between conditions (lmm, *p >* 0.282 for all pairwise comparisons). Flies from eggs associated with *Ac. thailandicus* developed faster than from axenic eggs or eggs associated with 11 out of the other 15 bacteria (lmm, *p* < 0.038 for these pairwise comparisons). (E, F) Fertility of G0 was assessed. Two males and one female that developed in the presence of different bacteria (G0) were placed per vial and flipped every other day for (E) or (F) days. Five couples were made per condition. Total number of emerged adults was analyzed. Flies associated with *Ac. thailandicus* are more fertile than axenic flies or flies associated with out of the other 15 bacteria (lmm, *p* < 0.018). (A, C, E) and (B, D, F) correspond to two experimental replicates. Correlation between developmental time and fertility is represented in Fig 7B. Each dot represents the total progeny of one female (A, B, E, F) and the size of the circle represents the mean number of adults that emerged per day (C, D). Statistical groups of significance for C, D, E, F are shown in S13 Fig.

**S13 Fig - Statistical groups of significance for developmental time and fertility of flies associated with different bacterial isolates.** Developmental time to adulthood (A) and fertility (B) of flies associated with different bacterial isolates from S12 Fig was analyzed with Tukey’s pairwise comparisons on the lmm estimates. Statistical groups of significance were generated with *cld* function in R. Groups with the same letter are not significantly different from each other.

**S14 Fig *-Ac. thailandicus* is beneficial for *D. melanogaster* in a natural food source.** (A, B) Thirty axenic *w^1118^ iso* eggs were placed in vials containing sterilized fig homogenate. *Ac. thailandicus* or sterile culture media were added on the top of the eggs. Four to six vials were used per condition. Total number of adults that emerged (A) and developmental time to adulthood (B) was determined. More eggs inoculated with *Ac. thailandicus* developed to adulthood and faster than axenic eggs (lmm, *p* < 0.001 for both comparisons). (C) Fertility of flies developed in fig homogenate with and without the addition of *Ac. thailandicus*. Two males and one female were collected from G0 and placed per vial containing fig homogenate for days, with vials flipped every other day. The *Ac. thailandicus* condition has ten replicates but only one from axenic eggs was possible to perform. Adults from eggs inoculated with *Ac. thailandicus* were more fertile than axenic adults (lmm, *p* = 0.003). (D, E) Fifty axenic *w^1118^ iso* eggs were placed in vials containing freshly collected non-sterile figs. *Ac. thailandicus* culture or sterile media (Control) was added on the top of the eggs. The total number of adults that emerged (D) and their developmental time to adulthood (E) was analyzed. Ten vials were analyzed per condition. There were more adults emerging from vials inoculated with *Ac. thailandicus* (lmm, *p* = 0.010). Developmental time to adulthood was not significantly different in this experimental replicate but faster in eggs inoculated with *Ac. thailandicus* in the other replicate represented on Fig 7J (lmm, *p* = 0.557 and *p* < 0.001, respectively). Statistical analyses were performed together with replicate experiments shown in Fig 7D-J.

**S1 Text - Sequences of the full 16S rRNA gene of the bacteria used in the phenotypic assays.** Sequence obtained by amplifying the gene with the primers 27F and 1495r. Code corresponds to code of laboratory isolate. It is also shown results of analysis on Greengenes, and of the BLAST analysis against the NCBI 16S ribosomal RNA sequences (Bacteria and Archaea) database.

**S2 Text – R script for data analysis.** Text is in R Markdown format.

**S1 Data - Bacterial levels from *w^1118^ iso* before and after 10 days in the same vial.** Bacterial numbers calculated per gut from each culture media used (BHI, LB, MRS, Mannitol or Liver) at day or day of the protocol. Data for Fig 1A and S1A Fig.

**S2 Data - Bacterial levels from *w^1118^ iso* before and after 10 days of being flipped to new vials twice a day.** Bacterial numbers calculated per gut from each culture media used (BHI, LB, MRS, Mannitol or Liver) at day 0 or day 10 of the protocol. Data for Fig 1B and S1A Fig.

**S3 Data - Relative 16S rRNA copy number *w^1118^ iso* before and 10 days after being flipped to new vials twice a day.** Data for Fig 1C and S1B Fig.

**S4 Data - Bacterial levels on the food after inoculating the food with flies.** Bacterial numbers calculated per food vial from each culture media used (BHI, LB, MRS, Mannitol or Liver) one or ten days after placing one fly. Data for Fig 1B and S1A Fig.

**S5 Data - Bacterial levels from wild caught flies before and 10 days after being flipped to new vials twice a day.** Bacterial numbers calculated per gut from each culture media used (BHI, LB, MRS, Mannitol or Liver) at day 0 or day 10 of the protocol. Data for Fig 1E, S1C and S1D Fig.

**S6 Data - Database for bacterial isolates that were sequenced and classified.** Source - sample origin. Day - day of the stability protocol. Media and Dilution - culture media and respective dilution from where colonies were isolated. Fly –gut sample number. Cfu_plate and Cfu_gut - Number of colonies analyzed in the plate and calculated per gut. morphotype - morphological type for one medium and one dilution. There is no correspondence with the same morphotype number in different media or flies. Bact_Code - code of laboratory isolate. greengenes_tax_string - list of taxonomic assignment according to Greengenes taxonomy. greengenes_prokMSA_id - identifier for the nearest neighbor sequence in the Greengenes database. greengenes_Simrank_id - percent of 7mers shared between the query sequence and the nearesr neighbor sequence. greengenes_DNAML_id - identity between the query and the nearest neighbor sequences. greengenes_DNAML_columns - number of bases compared between the query and the nearest neighbor sequences. sequence – 16S rRNA gene partial sequence. Fly_ID - Concatenation of Source, Day and Fly information. Unique_morpho - concatenation of Source, Day, Fly, Media and morphotype information. Data for Fig 2, Fig 3, S2, S3 and S4 Fig.

**S7 Data - Stability of *w^1118^ iso* monoassociated with different bacteria before and after being exposed to the stability protocol in vials and in cages.** Data for Fig 4B-E, S5A- S5I Fig.

**S8 Data - Stability of *Ac. thailandicus* in monoassociated *w^1118^ iso* with in different gut regions.** Data for Fig 4G, H and S5J, S5K Fig.

**S9 Data - Proliferation of different *Acetobacter* species and *Leuconostoc* in *w^1118^ iso*.** Data for Fig 4J-M and S6A-S6F Fig.

**S10 Data - Proliferation of *Lactobacillus* species in *w^1118^ iso*.** Data for Fig 4N, and S6H, S6J Fig.

**S11 Data *- Acetobacter* growth on the food.** Data for S7 Fig.

**S12 Data - Proliferation of *Ac. thailandicus* in *w^1118^ iso* during 24h in bottles with chaser GF flies.** Data for S8A, S8B Fig.

**S13 Data - Proliferation of *Ac. thailandicus* in *D. melanogaster* and *D. simulans*.** Data for Fig 5 and S8C-S8E Fig.

**S14 Data - Developmental time to pupariation and to adulthood of *w^1118^ iso* monoassociated with *Ac. thailandicus, Ac*. OTU2753 and axenic.** Column D-P correspond to number of new pupae or adults on the days 6-18 after egg laying. Data for S9 Fig.

**S15 Data - Colonization of *Ac. thailandicus* and *Ac*. OTU2753 in *w^1118^ iso* males and females at 0 days and 10 days after inoculation with the bacteria and being exposed to the stability protocol.** nc – Growth of bacteria in lowest dilution plate is too high to determine precisely CFUs. This data is represented as “above 10^5^ CFU/gut” in figures. Data for Fig 6B and S10A Fig.

**S16 Data - Transmission of *Ac. thailandicus* to food.** Column D-H correspond to assessment of bacteria in the food on days 1, 3, 5, 7, and 9 of the experiment. No data – no data collected. nc – Growth of bacteria in lowest dilution plate is too high to determine precisely CFUs. Data for Fig 6C and S10B Fig.

**S17 Data - Number of eggs laid by *w^1118^ iso* inoculated with *Ac. thailandicus, Ac*. OTU2753 or control, over the 10 days in cages.** Column D-M correspond to number of eggs on days 1-10 of experiment. Data for Fig 6D and S10C Fig.

**S18 Data - Developmental time of progeny from *w^1118^ iso* inoculated with *Ac. thailandicus, Ac*. OTU2753 or control, over the 10 days in cages.** Columns E-O correspond to number of new emerged adults on vials corresponding to days 10-20 after egg laying. Data for Fig 6E, F and S10D, S10E Fig.

**S19 Data - Fertility from progeny from *w^1118^ iso* inoculated with *Ac. thailandicus, Ac*. OTU2753 or control.** cagepair - cage from where the pairs were collected. daypair – pairs were collected from bottles of day 9 or 10 of the experiment. vialday – vial date; pairs were placed in new food vials every other day until day 8. Columns H-V correspond to number of new emerged adults on days 8-after egg laying. Data for Fig 6G and S10F Fig.

**S20 Data - Developmental time to pupariation, adulthood and respective total number of progeny from one or both parents monoassociated with *Ac. thailandicus*.** Conditions used were: 53F + 53M – both parents associated with *Ac. thailandicus*. GFF + GFM – both parents axenic. 53F + GFM – female with *Ac. thailandicus*, male axenic. GFF + 53M – female axenic, male with *Ac. thailandicus*. GFF + GFM + Bact – both parents axenic and *Ac. thailandicus* added. vialday – vial date; pairs were placed in new food vials every other day until day 8. Columns F-W correspond to number of new pupae or emerged adults on days 5-22 after egg laying. Data for Fig 6H and S11 Fig.

**S21 Data - Developmental time to adulthood from *w^1118^ iso* associated with different bacterial isolates.** Columns E-O correspond to number of new emerged adults on days 9-19 after egg laying. Data for Fig 7B, S12A-S12D Fig and S13B Fig.

**S22 Data - Fertility of *w^1118^ iso* developed with different bacterial isolates.** Data for Fig 7B, S12E, S12F Fig and S13A Fig.

**S23 Data - Developmental time of axenic *w^1118^ iso* or associated with *Ac. thailandicus* in sterilized fig food.** Columns D-AD correspond to number of new emerged adults on days 9-after egg laying. Data for Fig 7D, E and S12A-S12D Fig.

**S24 Data - Fertility of axenic *w^1118^ iso* or associated with *Ac. thailandicus* in fig food.** Data for Fig 7G and S13C Fig.

**S25 Data - Developmental time of *w^1118^ iso* with and without the addition of *Ac. thailandicus* in freshly collected figs.** Columns E-U correspond to number of new emerged adults on 1868 days 9-25 after egg laying. Data for Fig7I, J and S14D, S14E Fig.

## References

1. McFall-Ngai M, Hadfield MG, Bosch TCG, Carey HV, Domazet-Lošo T, Douglas AE, et al. Animals in a bacterial world, a new imperative for the life sciences. Proc Natl Acad Sci USA. 2013;110: 3229–3236. doi:10.1073/pnas.1218525110

2. Sasaki T, Ishikawa H. Production of Essential Amino Acids from Glutamate by Mycetocyte Symbionts of the Pea Aphid, Acyrthosiphon pisum. J Insect Physiol. 1995;41: 41–46.

3. Douglas AE. Reproductive failure and the free amino acid pools in pea aphids (Acyrthosiphon pisum) lacking symbiotic bacteria. J Insect Physiol. 1996;42: 247–255. doi:10.1016/0022-1910(95)00105-0

4. Akman L, Yamashita A, Watanabe H, Oshima K, Shiba T, Hattori M, et al. Genome sequence of the endocellular obligate symbiont of tsetse flies, Wigglesworthia glossinidia. Nat Genet. 2002;32: 402–407. doi:10.1038/ng986

5. Pais R, Lohs C, Wu Y, Wang J, Aksoy S. The obligate mutualist Wigglesworthia glossinidia influences reproduction, digestion, and immunity processes of its host, the tsetse fly. Appl Environ Microbiol. 2008;74: 5965–5974. doi:10.1128/AEM.00741-08

6. Hosokawa T, Koga R, Kikuchi Y, Meng X-Y, Fukatsu T. Wolbachia as a bacteriocyte-associated nutritional mutualist. Proc Natl Acad Sci USA. 2010;107: 769–774. doi:10.1073/pnas.0911476107

7. Qin J, Li R, Raes J, Arumugam M, Burgdorf KS, Manichanh C, et al. A human gut microbial gene catalogue established by metagenomic sequencing. Nature. 2010;464: 59–65. doi:10.1038/nature08821

8. Spor A, Koren O, Ley R. Unravelling the effects of the environment and host genotype on the gut microbiome. Nature Rev Microbiology 2011;9: 279–290. doi:10.1038/nrmicro2540

9. Faith JJ, Mcnulty NP, Rey FE, Gordon JI. Predicting a Human Gut Microbiota’s Response to Diet in Gnotobiotic Mice. Science. 2011;333: 101–104. doi:10.1186/1471-2164-10-161

10. Kurokawa K, Itoh T, Kuwahara T, Oshima K, Toh H, Toyoda A, et al. Comparative Metagenomics Revealed Commonly Enriched Gene Sets in Human Gut Microbiomes. DNA Res. 2007;14: 169–181. doi:10.1093/dnares/dsm018

11. Arumugam M, Raes J, Pelletier E, Le Paslier D, Yamada T, Mende DR, et al. Enterotypes of the human gut microbiome. Nature. 2011;473: 174–180. doi:10.1038/nature09944

12. Martino ME, Ma D, Leulier F. Microbial influence on Drosophila biology. Curr Opin Microbiol. 2017;38: 165–170. doi:10.1016/j.mib.2017.06.004

13. Broderick NA, Lemaitre B. Gut-associated microbes of Drosophila melanogaster. Gut Microbes. 2012;3: 307–321. doi:10.4161/gmic.19896

14. Baumberg JP. A nutriotional study of insects, with special reference to microorganisms and their substrata. Journal of Experimental Zoology Part A: Ecological Genetics and Physiology. 1919;28: 1–81.

15. Storelli G, Defaye A, Erkosar B, Hols P, Royet J, Leulier F. Lactobacillus plantarum promotes Drosophila systemic growth by modulating hormonal signals through TOR-dependent nutrient sensing. Cell Metab. 2011; 14: 403–414. doi:10.1016/j.cmet.2011.07.012

16. Shin SC, Kim S-H, You H, Kim B, Kim AC, Lee K-A, et al. Drosophila microbiome modulates host developmental and metabolic homeostasis via insulin signaling. Science. 2011;334: 670–674. doi:10.1126/science.1212782

17. Buchon N, Broderick NA, Lemaitre B. Gut homeostasis in a microbial world: insights from Drosophila melanogaster. Nature Rev Microbiology. 2013; 11: 615–626. doi:10.1038/nrmicro3074

18. Dantoft W, Lundin D, Esfahani SS, Engström Y. The POU/Oct Transcription Factor Pdm1/nub Is Necessary for a Beneficial Gut Microbiota and Normal Lifespan of Drosophila. J Innate Immun. 2016;8: 412–426. doi:10.1159/000446368

19. Yamada R, Deshpande SA, Bruce KD, Mak EM, Ja WW. Microbes Promote Amino Acid Harvest to Rescue Undernutrition in Drosophila. Cell Reports. 2015;0. doi:10.1016/j.celrep.2015.01.018

20. Blum JE, Fischer CN, Miles J, Handelsman J. Frequent replenishment sustains the beneficial microbiome of Drosophila melanogaster. MBio. 2013;4: e00860–13. doi:10.1128/mBio.00860-13

21. Glittenberg M, Kounatidis I, Christensen D, Kostov M, Kimber S, Roberts I, et al. Pathogen and host factors are needed to provoke a systemic host response to gastrointestinal infection of Drosophila larvae by Candida albicans. Disease models & mechanisms. 2011. doi:10.1242/dmm.006627

22. Ryu J-H, Kim S-H, Lee H-Y, Bai JY, Nam Y-D, Bae J-W, et al. Innate immune homeostasis by the homeobox gene caudal and commensal-gut mutualism in Drosophila. Science. 2008;319: 777–782. doi:10.1126/science.1149357

23. Wong AC-N, Wang Q-P, Morimoto J, Senior AM, Lihoreau M, Neely GG, et al. Gut Microbiota Modifies Olfactory-Guided Microbial Preferences and Foraging Decisions in Drosophila. Curr Biol 2017;27: 2397–2404.e4. doi:10.1016/j.cub.2017.07.022

24. Ren C, Webster P, Finkel SE, Tower J. Increased internal and external bacterial load during Drosophila aging without life-span trade-off. Cell Metab. 2007;6: 144–152. doi:10.1016/j.cmet.2007.06.006

25. Chandler JA, Morgan Lang J, Bhatnagar S, Eisen JA, Kopp A. Bacterial communities of diverse Drosophila species: ecological context of a host-microbe model system. PLoS Genet. 2011;7: e1002272. doi:10.1371/journal.pgen.1002272

26. Wong CNA, Ng P, Douglas AE. Low-diversity bacterial community in the gut of the fruitfly Drosophila melanogaster. Environ Microbiol. 2011; 13: 1889–1900. doi:10.1111/j.1462-2920.2011.02511.x

27. Broderick NA, Buchon N, Lemaitre B. Microbiota-induced changes in drosophila melanogaster host gene expression and gut morphology. MBio. 2014;5: e01117–14. doi:10.1128/mBio.01117-14

28. Staubach F, Baines JF, Künzel S, Bik EM, Petrov DA. Host Species and Environmental Effects on Bacterial Communities Associated with Drosophila in the Laboratory and in the Natural Environment. PLoS ONE. 2013;8: e70749. doi:10.1371/journal.pone.0070749

29. Martinson VG, Carpinteyro-Ponce J, Moran NA, Markow TA. A Distinctive and Host-Restricted Gut Microbiota in Populations of a Cactophilic Drosophila Species. Appl Environ Microbiol. 2017;83. doi:10.1128/AEM.01551-17

30. Martinson VG, Douglas AE, Jaenike J. Community structure of the gut microbiota in sympatric species of wild Drosophila. Ecology Letters. 2017;26: 32. doi:10.1111/ele.12761

31. Obadia B, Güvener ZT, Zhang V, Ceja-Navarro JA, Brodie EL, Ja WW, et al. Probabilistic Invasion Underlies Natural Gut Microbiome Stability. Curr Biol. 2017;27: 1999–2006.e8. doi:10.1016/j.cub.2017.05.034

32. Chrostek E, Marialva MSP, Esteves SS, Weinert LA, Martinez J, Jiggins FM, et al. Wolbachia Variants Induce Differential Protection to Viruses in Drosophila melanogaster: A Phenotypic and Phylogenomic Analysis. PLoS Genet. 2013;9: e1003896. doi:10.1371/journal.pgen.1003896

33. Ryder E, Blows F, Ashburner M, Bautista-Llacer R, Coulson D, Drummond J, et al. The DrosDel collection: a set of P-element insertions for generating custom chromosomal aberrations in Drosophila melanogaster. Genetics. 2004;167: 797–813. doi:10.1534/genetics.104.026658

34. DeSantis TZ, Hugenholtz P, Larsen N, Rojas M, Brodie EL, Keller K, et al. Greengenes, a chimera-checked 16S rRNA gene database and workbench compatible with ARB. Appl Environ Microbiol. 2006;72: 5069–5072. doi:10.1128/AEM.03006-05

35. NCBI Resource Coordinators. Database Resources of the National Center for Biotechnology Information. Nucleic Acids Res. 2017;45: D12–D17. doi:10.1093/nar/gkw1071

36. Pitiwittayakul N, Yukphan P, Chaipitakchonlatarn W, Yamada Y, Theeragool G. Acetobacter thailandicus sp. nov., for a strain isolated in Thailand. Ann Microbiol. 2015;65: 1855–1863. doi:10.1007/s13213-014-1024-7

37. Tefit M, Leulier F. Lactobacillus plantarum favors the early emergence of fit and fertile adult Drosophila upon chronic undernutrition. J Exp Biol. 2017;220: 900-907. doi:10.1242/jeb.151522

38. Storelli G, Strigini M, Grenier T, Bozonnet L, Schwarzer M, Daniel C, et al. Drosophila Perpetuates Nutritional Mutualism by Promoting the Fitness of Its Intestinal Symbiont Lactobacillus plantarum. Cell Metab. 2017. doi:10.1016/j.cmet.2017.11.011

39. Matos RC, Schwarzer M, Gervais H, Courtin P, Joncour P, Gillet B, et al. d-alanine esterification of teichoic acids contributes to Lactobacillus plantarum-mediated Drosophila growth promotion during chronic undernutrition. Nat Microbiol. 2017;2: 1635–1647. doi:10.1038/s41564-017-0038-x

40. Markow TA, O’Grady PM. Drosophila : a guide to species identification and use. Associated Press; 2006.

41. Corby-Harris V, Pontaroli AC, Shimkets LJ, Bennetzen JL, Habel KE, Promislow DEL. Geographical Distribution and Diversity of Bacteria Associated with Natural Populations of Drosophila melanogaster. 2007;73: 3470. doi:10.1128/AEM.02120-06

42. Cox CR, Gilmore MS. Native microbial colonization of Drosophila melanogaster and its use as a model of Enterococcus faecalis pathogenesis. Infect Immun. 2007;75: 1565–1576. doi:10.1128/IAI.01496-06

43. Wong AC-N, Chaston JM, Douglas AE. The inconstant gut microbiota of Drosophila species revealed by 16S rRNA gene analysis. The ISME Journal. 2013. doi:10.1038/ismej.2013.86

44. Wong AC-N, Luo Y, Jing X, Franzenburg S, Bost A, Douglas AE. The Host as Driver of the Microbiota in the Gut and External Environment of Drosophila melanogaster. Appl Environ Microbiol. 2015;81:6232-6240. doi:10.1128/AEM.01442-15

45. Clark RI, Salazar A, Yamada R, Fitz-Gibbon S, Morselli M, Alcaraz J, et al. Distinct Shifts in Microbiota Composition during Drosophila Aging Impair Intestinal Function and Drive Mortality. Cell Reports. 2015;12: 1656–67. doi:10.1016/j.celrep.2015.08.004

46. Wang Y, Staubach F. Individual Variation Of Natural D. melanogaster Associated Bacterial Communities. FEMS Microbiol Lett. 2018. doi:10.1101/126912

47. Miller A. The internal anatomy and histology of the imago of Drosophila melanogaster. In: Demerec M, editor. Biology of Drosophila. Cold Spring Harbor Laboratory Press; 1994.

48. Randal Bollinger R, Barbas AS, Bush EL, Lin SS, Parker W. Biofilms in the large bowel suggest an apparent function of the human vermiform appendix. J Gen Virol. 2007;249: 826–831. doi:10.1016/j.jtbi.2007.08.032

49. Brüssow H. How stable is the human gut microbiota? And why this question matters. Environ Microbiol. 2016;18: 2779–2783. doi:10.1111/1462-2920.13473

50. Hegedus D, Erlandson M, Gillott C, Toprak U. New insights into peritrophic matrix synthesis, architecture, and function. Annu Rev Entomol. 2009;54: 285–302. doi:10.1146/annurev.ento.54.110807.090559

51. Lemaitre B, Miguel-Aliaga I. The digestive tract of Drosophila melanogaster. Annu Rev Genet. 2013;47: 377–404. doi:10.1146/annurev-genet-111212-133343

52. Lanan MC, Rodrigues PAP, Agellon A, Jansma P, Wheeler DE. A bacterial filter protects and structures the gut microbiome of an insect. The ISME Journal. 2016;10: 1866–1876. doi:10.1038/ismej.2015.264

53. Hao Z, Kasumba I, Aksoy S. Proventriculus (cardia) plays a crucial role in immunity in tsetse fly (Diptera: Glossinidiae). Insect Biochem Mol Biol. 2003;33: 1155–1164. doi:10.1016/j.ibmb.2003.07.001

54. Liehl P, Blight M, Vodovar N, Boccard F, Lemaitre B. Prevalence of local immune response against oral infection in a Drosophila/Pseudomonas infection model. PLoS Pathog. 2006;2: e56. doi:10.1371/journal.ppat.0020056

55. Li H, Qi Y, Jasper H. Preventing Age-Related Decline of Gut Compartmentalization Limits Microbiota Dysbiosis and Extends Lifespan. Cell Host Microbe. 2016; 19: 240–253. doi:10.1016/j.chom.2016.01.008

56. Mulcahy H, Sibley CD, Surette MG, Lewenza S. Drosophila melanogaster as an animal model for the study of Pseudomonas aeruginosa biofilm infections in vivo. PLoS Pathog. 2011;7: e1002299. doi:10.1371/journal.ppat.1002299

57. Giglioli I. Insects and yeasts. Nature. 1897;56: 575–577.

58. Purdy AE, Watnick PI. Spatially selective colonization of the arthropod intestine through activation of Vibrio cholerae biofilm formation. Proceedings of the National Academy of Sciences. 2011;108: 19737–19742. doi:10.1073/pnas.1111530108

59. Jarrett CO, Deak E, Isherwood KE, Oyston PC, Fischer ER, Whitney AR, et al. Transmission of Yersinia pestis from an infectious biofilm in the flea vector. J Infect Dis. 2004; 190: 783–792. doi:10.1086/422695

60. Powell JE, Martinson VG, Urban-Mead K, Moran NA. Routes of Acquisition of the Gut Microbiota of the Honey Bee Apis mellifera. Appl Environ Microbiol. 2014;80: 7378–7387. doi:10.1128/AEM.01861-14

61. Ohbayashi T, Takeshita K, Kitagawa W, Nikoh N, Koga R, Meng X-Y, et al. Insect’s intestinal organ for symbiont sorting. Proceedings of the National Academy of Sciences. 2015;: 201511454. doi:10.1073/pnas.1511454112

62. Dillon RJ, Dillon VM. The gut bacteria of insects: nonpathogenic interactions. Annu Rev Entomol. 2004;49: 71–92. doi:10.1146/annurev.ento.49.061802.123416

63. Mead LJ, Khachatourians GG, Jones GA. Microbial Ecology of the Gut in Laboratory Stocks of the Migratory Grasshopper, Melanoplus sanguinipes (Fab.) (Orthoptera: Acrididae). Appl Environ Microbiol. 1988;54: 1174–1181.

64. Garcia JR, Gerardo NM. The symbiont side of symbiosis: do microbes really benefit? Front Microbiol. 2014;5: 510. doi:10.3389/fmicb.2014.00510

65. Mushegian AA, Ebert D. Rethinking “mutualism” in diverse host-symbiont communities. Bioessays. 2016;38: 100–108. doi:10.1002/bies.201500074

66. Bakula M. The persistence of a microbial flora during postembryogenesis of Drosophila melanogaster. Journal of Invertebrate Pathology. 1969; 14: 365–374.

67. Chaston JM, Newell PD, Douglas AE. Metagenome-Wide Association of Microbial Determinants of Host Phenotype in Drosophila melanogaster. MBio. 2014;5: e01631–14–e01631–14. doi:10.1128/mBio.01631-14

68. Wong AC-N, Dobson AJ, Douglas AE. Gut microbiota dictates the metabolic response of Drosophila to diet. J Exp Biol. 2014;217: jeb.101725–1901. doi:10.1242/jeb.101725

69. Soen Y. Delayed development induced by toxicity to the host can be inherited by a bacterial-dependent, transgenerational effect. Frontiers in Genetics. 2014;5: 1–14. doi:10.3389/fgene.2014.00027/abstract

70. Elgart M, Stern S, Salton O, Gnainsky Y, Heifetz Y, Soen Y. Impact of gut microbiota on the fly’s germ line. Nat Comms. 2016;7: 11280. doi:10.1038/ncomms11280

71. Sang JH. The Nutriotional Requirements of Drosophila. In: Ashburner M, Wright T, editors. The Genetics and Biology of Drosophila. 1978. pp. 159–192.

72. David J. A new medium for rearing Drosophila in axenic conditions. Drosophila Information Service. 1962;36: 128.

73. Fridmann-Sirkis Y, Stern S, Elgart M, Galili M, Zeisel A, Shental N, et al. Delayed development induced by toxicity to the host can be inherited by a bacterial-dependent, transgenerational effect. Front Genet. 2014;5: 27. doi:10.3389/fgene.2014.00027

74. Erkosar B, Storelli G, Mitchell M, Bozonnet L, Bozonnet N, Leulier F. Pathogen Virulence Impedes Mutualist-Mediated Enhancement of Host Juvenile Growth via Inhibition of Protein Digestion. Cell Host Microbe. 2015;18: 445–455. doi:10.1016/j.chom.2015.09.001

75. van den Bosch TJM, Welte CU. Detoxifying symbionts in agriculturally important pest insects. Microbial Biotechnology. 2017; 10: 531–540. doi:10.1111/1751-7915.12483

76. Leitão-Gonçalves R, Carvalho-Santos Z, Francisco AP, Fioreze GT, Anjos M, Baltazar C, et al. Commensal bacteria and essential amino acids control food choice behavior and reproduction. PLOS Biol. 2017; 15: e2000862. doi:10.1371/journal.pbio.2000862

77. Mueller UG, Gerardo NM, Aanen DK, Six DL, Schultz TR. The Evolution of Agriculture in Insects. Annu Rev Ecol Evol S. 2005;36: 563–595. doi:10.1146/annurev.ecolsys.36.102003.152626

78. Venu I, Durisko Z, Xu J, Dukas R. Social attraction mediated by fruit flies’ microbiome. Journal of Experimental Biology. 2014;217: 1346–1352. doi:10.1242/jeb.099648

79. Fuyama Y. Behavior genetics of olfactory responses in Drosophila. I. Olfactometry and strain differences in Drosophila melanogaster. Behav Genet. 1976;6: 407–420.

80. Fischer CN, Trautman EP, Crawford JM, Stabb EV, Handelsman J, Broderick NA. Metabolite exchange between microbiome members produces compounds that influence Drosophila behavior. Elife. 2017;6: e18855. doi:10.7554/eLife.18855

81. Mansourian S, Stensmyr MC. The chemical ecology of the fly. Current Opinion in Neurobiology. 2015;34: 95–102. doi:10.1016/j.conb.2015.02.006

82. Crotti E, Rizzi A, Chouaia B, Ricci I, Favia G, Alma A, et al. Acetic acid bacteria, newly emerging symbionts of insects. Appl Environ Microbiol. 2010;76: 6963–6970. doi:10.1128/AEM.01336-10

83. Kounatidis I, Crotti E, Sapountzis P, Sacchi L, Rizzi A, Chouaia B, et al. Acetobacter tropicalis Is a Major Symbiont of the Olive Fruit Fly (Bactrocera oleae). Appl Environ Microbiol. 2009;75: 3281–3288. doi:10.1128/AEM.02933-08

84. Mazzetto F, Gonella E, Crotti E, Vacchini V, Syrpas M, Pontini M, et al. Olfactory attraction of Drosophila suzukii by symbiotic acetic acid bacteria. J Pest Sci. 2016;89: 783–792. doi:10.1007/s10340-016-0754-7

85. Chouaia B, Gaiarsa S, Crotti E, Comandatore F, Degli Esposti M, Ricci I, et al. Acetic acid bacteria genomes reveal functional traits for adaptation to life in insect guts. Genome Biol Evol. 3rd ed. 2014;6: 912–920. doi:10.1093/gbe/evu062

86. Engel P, Moran NA. The gut microbiota of insects - diversity in structure and function. FEMS Microbiol Rev. 2013;37: 699–735. doi:10.1111/1574-6976.12025

87. Dworkin M, Falkow S, Rosenberg E, Schleifer K-H, Stackebrandt E, editors. The Prokaryotes. Springer New York; 2006. doi:10.1007/0-387-30745-1

88. Crotti E, Damiani C, Pajoro M, Gonella E, Rizzi A, Ricci I, et al. Asaia, a versatile acetic acid bacterial symbiont, capable of cross-colonizing insects of phylogenetically distant genera and orders. Environ Microbiol. 2009;11: 3252–3264. doi:10.1111/j.1462-2920.2009.02048.x

89. Sharon G, Segal D, Ringo JM, Hefetz A, Zilber-Rosenberg I, Rosenberg E. Commensal bacteria play a role in mating preference of Drosophila melanogaster. Proc Natl Acad Sci USA. 2010; 107: 20051–20056. doi:10.1073/pnas.1009906107

90. Powell E, Ratnayeke N, Moran NA. Strain diversity and host specificity in a specialized gut symbiont of honeybees and bumblebees. Mol Ecol. 2016;25: 4461–4471. doi:10.1111/mec.13787

91. Moeller AH, Caro-Quintero A, Mjungu D, Georgiev AV, Lonsdorf EV, Muller MN, et al. Cospeciation of gut microbiota with hominids. Science. 2016;353: 380–382. doi:10.1126/science.aaf3951

92. Brooks AW, Kohl KD, Brucker RM, van Opstal EJ, Bordenstein SR. Phylosymbiosis: Relationships and Functional Effects of Microbial Communities across Host Evolutionary History. PLOS Biol. 2016;14: e2000225. doi:10.1371/journal.pbio.2000225

93. Finlay BB, McFadden G. Anti-immunology: evasion of the host immune system by bacterial and viral pathogens. Cell. 2006;124: 767–782. doi:10.1016/j.cell.2006.01.034

94. Broderick NA. Friend, foe or food? Recognition and the role of antimicrobial peptides in gut immunity and Drosophila-microbe interactions. Philos Trans R Soc Lond, B, Biol Sci. 2016;371: 20150295–10. doi:10.1098/rstb.2015.0295

95. Pang X, Xiao X, Liu Y, Zhang R, Liu J, Liu Q, et al. Mosquito C-type lectins maintain gut microbiome homeostasis. Nat Microbiol. 2016;: 16023. doi:10.1038/nmicrobiol.2016.23

96. Sackton TB, Lazzaro BP, Schlenke TA, Evans JD, Hultmark D, Clark AG. Dynamic evolution of the innate immune system in Drosophila. Nat Genet. 2007;39: 1461–1468. doi:10.1038/ng.2007.60

97. Obbard DJ, Welch JJ, Kim K-W, Jiggins FM. Quantifying adaptive evolution in the Drosophila immune system. PLoS Genet. 2009;5: e1000698. doi:10.1371/journal.pgen.1000698

98. Lazzaro BP. Natural selection on the Drosophila antimicrobial immune system. Curr Opin Microbiol. 2008;11: 284–289. doi:10.1016/j.mib.2008.05.001

99. Erkosar B, Leulier F. Transient adult microbiota, gut homeostasis and longevity: novel insights from the Drosophila model. FEBS Lett. 2014;588: 4250–4257. doi:10.1016/j.febslet.2014.06.041

100. Winans NJ, Walter A, Chouaia B, Chaston JM, Douglas AE, Newell PD. A genomic investigation of ecological differentiation between free-living and Drosophila-associated bacteria. Mol Ecol. 2017;26: 4536–4550. doi:10.1111/mec.14232

101. Samuel BS, Rowedder H, Braendle C, Félix M-A, Ruvkun G. Caenorhabditis elegans responses to bacteria from its natural habitats. Proc Natl Acad Sci USA. 2016; 113: E3941–9. doi:10.1073/pnas.1607183113

102. Dirksen P, Marsh SA, Braker I, Heitland N, Wagner S, Nakad R, et al. The native microbiome of the nematode Caenorhabditis elegans: gateway to a new host-microbiome model. BMC Biol. 2016; 14: 38. doi:10.1186/s12915-016-0258-1

103. Rosshart SP, Vassallo BG, Angeletti D, Hutchinson DS, Morgan AP, Takeda K, et al. Wild Mouse Gut Microbiota Promotes Host Fitness and Improves Disease Resistance. Cell. 2017; 171: 1015–1028.e13. doi:10.1016/j.cell.2017.09.016

104. Cowles CE, Goodrich Blair H. The Xenorhabdus nematophila nilABC genes confer the ability of Xenorhabdus spp. to colonize Steinernema carpocapsae nematodes. J Bacteriol. 2008; 190: 4121–4128. doi:10.1128/JB.00123-08

105. Frese SA, MacKenzie DA, Peterson DA, Schmaltz R, Fangman T, Zhou Y, et al. Molecular characterization of host-specific biofilm formation in a vertebrate gut symbiont. PLoS Genet. 2013;9: e1004057. doi:10.1371/journal.pgen.1004057

106. Yuval B, Ben Ami E, Behar A. The Mediterranean fruit fly and its bacteria–potential for improving sterile insect technique operations. Journal of Applied Entomology. 2013; 137: 39–42. doi:10.1111/j.1439-0418.2010.01555.x

107. Dong Y, Manfredini F, Dimopoulos G. Implication of the mosquito midgut microbiota in the defense against malaria parasites. PLoS Pathog. 2009;5: e1000423. doi:10.1371/journal.ppat.1000423

108. Cirimotich CM, Dong Y, Clayton AM, Sandiford SL, Souza-Neto JA, Mulenga M, et al. Natural microbe-mediated refractoriness to Plasmodium infection in Anopheles gambiae. Science. 2011;332: 855–858. doi:10.1126/science.1201618

109. Caporaso JG, Bittinger K, Bushman FD, DeSantis TZ, Andersen GL, Knight R. PyNAST: a flexible tool for aligning sequences to a template alignment. Bioinformatics. 2010;26: 266–267. doi:10.1093/bioinformatics/btp636

110. Dridi B, Henry M, Khéchine El A, Raoult D, Drancourt M. High prevalence of Methanobrevibacter smithii and Methanosphaera stadtmanae detected in the human gut using an improved DNA detection protocol. PLoS ONE. 2009;4: e7063. doi:10.1371/journal.pone.0007063

111. Pfaffl MW. A new mathematical model for relative quantification in real-time RT-PCR. Nucleic Acids Res. 2001;29: e45.

112. Team RC. R: A language and environment for statistical computing [Internet]. R Foundation for Statistical Computing, Vienna, Austria; 2012. Available: URL http://www.R-project.org/

113. Wickham H. ggplot2 - Elegant Graphics for Data Analysis. Springer. 2009

114. Bates D, Mächler M, Bolker B, Walker S. Fitting Linear Mixed-Effects Models Using lme4. J Stat Soft. 2015;67: 1–48. doi:10.18637/jss.v067.i01

115. Kuznetsova A, Brockhoff PB, Christensen RHB. Tests in Linear Mixed Effects Models. Comprehensive R Archive Network (CRAN); 2016. Available: https://CRAN.R-project.org/package=lmerTest

116. Lenth R. Least-squares Means: The R Package lsmeans. J Stat Soft. 2016;69. doi:10.18637/jss.v069.i01

117. Hothom T, Bretz F, Westfall P. Simultaneous inference in general parametric models. Biom J. 2008;50: 346–363. doi:10.1002/bimj.200810425

